# GABA regulates resonance and spike rate encoding via a universal mechanism that underlies the modulation of action potential generation

**DOI:** 10.1101/206581

**Authors:** Christoph Kirst, Julian Ammer, Felix Felmy, Andreas Herz, Martin Stemmler

## Abstract

Different mechanisms for action potential (AP) generation exist that shape neuronal coding and network dynamics. The neuro-transmitter GABA regulates neuronal activity but its role in modulating AP dynamics itself is unclear. Here we show that GABA indeed changes the AP mechanism: it causes regularly firing hippocampal CA3 neurons to bistably switch between spiking and quiescence, converts graded discharge-to-current relationships to have abrupt onsets, and induces resonance. Modeling reveals that A-currents enable these GABA-induced transitions. Mathematically, we prove that this transition sequence originates from a single universal principle that generically underlies the modulation of AP dynamics in any conductance-based neuron model. Conductance clamp experiments in hippocampal and brainstem neurons show the same transitions, confirming the universal theory. In simulated networks, synaptically controlled AP dynamics, permits dynamic gating of signals and targeted synchronization of neuronal sub-ensembles. These results advance the systematic understanding of AP modulation and its role in neuronal and network function.

## Introduction

Neuronal action potentials (APs) are fundamental to communication and computation in the nervous system. How neurons decide to respond to inputs, code information, and participate in network dynamics depends on the precise dynamical mechanisms underlying the generation of APs (Rinzel & Ermentrout, 1989; Hansel et al., 1995; Ermentrout, 1996; Koch, 1997; Gutkin & Ermentrout, 1998; Robinson & Harsch, 2002; Fourcaud-Trocme et al., 2003; Gutkin et al., 2005; Izhikevich, 2010; Schleimer & Stemmler, 2009; Ermentrout & Terman, 2010; Hong et al., 2012; Ratté et al., 2013). AP dynamics are governed by a host of currents, which are subject to intrinsic cellular plasticity and numerous neuro-modulatory signals (Kaczmarek & Levitan, 1987; Getting, 1989; Golowasch et al., 1999; Turrigiano et al., 1994; Prescott et al., 2006; Stiefel et al., 2008; Prescott et al., 2008b; Bargmann, 2012; Marder, 2012; Cudmore & Turrigiano, 2015; Drion et al., 2015; Morozova et al., 2016). Given these multiple influences on AP generation, identifying general principles underlying the modulation of AP dynamics will aid in understanding the adaptive function of neurons and neuronal circuits.

The neuro-transmitter gamma-aminobutyric acid (GABA), prevalent throughout the nervous system, regulates neuronal activity (Cobb et al., 1995; Semyanov et al., 2004; Gulledge & Stuart, 2003; Mody & Pearce, 2004; Vida et al., 2006; Mann & Paulsen, 2007; Jeong & Gutkin, 2007; Olah et al., 2009; Deng et al., 2009; Brickley & Mody, 2012; Ammer et al., 2012; Pavlov et al., 2014), normalizes neuronal responses (Carandini & Heeger, 2012; Chance et al., 2002; Holt & Koch, 1997) and regulates neuronal rhythms (Mann & Paulsen, 2007; Buzsaki, 2006). However, whether GABA can temper the very mechanism of generating APs has not been reported.

The CA3 region in the hippocampus lends itself to testing this question, as it is subject to acute GABAergic inhibition (Andersen et al., 1964; Mann & Mody, 2010) that balances extensive recurrent excitation (Johston & Amaral, 2004), which may serve auto-associative functions (Marr, 1971; Cherubini & Miles, 2015) but also facilitates epileptic activity (Traub et al., 1987, 1992). The flow of excitation through the hippocampus is regimented by GABA pulses from interneurons (Hasselmo, 2006; Mann & Paulsen, 2007; Buzsaki, 2006): phasic responses to GABA_A_ on the milli-second time scale are complemented by slower responses from metabotropic GABA_B_ and extra-and peri-synaptic GABA_A_ receptors on the time scale of 500-2000 ms (Scanziani, 2000; Farrant & Nusser, 2005; Semyanov et al., 2004; Cherubini & Miles, 2015). In particular, the slower components may modulate the AP generation mechanism, with potential consequences for network function and in pathology.

Here we show that GABA controls AP generation dynamics and identify this effect as part of a general mechanisms that underlies the modulation of AP dynamics. In pyramidal hippocampal CA3 neurons, GABA forces a continuous relationship between AP frequency and input current (*f-I*-curve) to become discontinuous, it induces bistable dynamics in which spiking alternates with quiescence, and it causes non-resonant neurons to resonate in the range of known physiological rhythms. According to the classical categorization of neuronal firing patterns by Hodgkin (1948), the observed switch from continuous to discontinuous *f-I*-curves represents a switch from class-I to class-II excitability. Using dynamic clamp experiments, we were able to mimic the same entire sequence of transitions by increasing the leak conductance both in CA3 and also brain stem neurons, consistent with transitions in neuronal excitability in CA1 neurons (Prescott et al., 2008b). Drawing on the theory of how higher-order bifurcation points unfold in parameter spaces of nonlinear dynamical systems (Dumortier et al., 1991), we developed a mathematical approach that explains the entire sequence of transitions in AP generation dynamics. This sequence occurs universally in all regularly firing class-I excitable neurons, regardless of the particular expression patterns of their ionic currents. Moreover, changing any malleable neuronal property affecting the AP mechanism leads to the same set of transitions.

A-type conductances are known to delay APs and to move class-II neurons towards class-I excitability (Connor & Stevens, 1971; Ermentrout, 1996). Via modeling we here find the opposite effect, namely that the A-type conductance supports the role of GABA in inducing transitions from class-I to II and towards resonance. This mechanisms is distinct from another effect that generates class-II excitability at very high A-type conductances (Drion et al., 2015). Our generic mathematical theory captures these effects and resolves how the A-type currents differentially modulate AP generation.

Changing the AP dynamics has immediate consequences for network function, as, neuronal excitability class and resonances influence synchronization in neuronal networks (Hansel et al., 1995; Ermentrout, 1996; Izhikevich, 2010). GABA-induced modulation of AP generation adds another facet: using computational modeling, we demonstrate that the control of neuronal excitability permits recurrent networks to dynamically group sub-assemblies and via the induction of bi-stability controls the maximum number of synchronized neurons.

## Results

### GABA-ergic modulation AP dynamics and of resonance in hippocampal pyramidal cells

To test how tonic GABA-ergic inhibition affects the dynamical characteristics of AP generation, we puffed GABA locally onto hippocampal CA3 pyramidal cells in brain slices of Mongolian gerbil and measured the neurons’ response to different stimuli near AP threshold. All measured pyramidal neurons in the CA3 area showed regular spiking and no bursting, in agreement with earlier studies (Hemond et al., 2008; Migliore et al., 2010; Graves et al., 2012; Magraner et al., 2017). This allowed us to focus on the effect of GABA on the AP generation mechanism itself. In response to step currents prior to GABA application, a typical cell exhibited slow firing close to AP threshold (Fig. 1A), with nearly constant AP amplitudes and stereotyped trajectories in the phase plane spanned by the voltage *V* and its time derivative 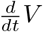. Puff application of GABA raised the current-threshold for spike generation (Fig. 1B), similar to what is observed in CA1 principal cells (Pavlov et al., 2009). In addition, the voltage amplitude of subsequent APs decayed in the presence of GABA (Fig. 1B). The *f-I*-curve, which measures the firing rate *f* as a function of the external input current *I_e_*, also altered its shape (Fig. 1C) by changing from a continuously rising curve (AP onset frequency *f*_*θ*_ < 1Hz) to one that has a discontinuous jump at AP onset (*f*_*θ*_ ≈ 10Hz). All CA3 neurons with a continuous *f-I*-curve (*n* = 6) showed this switch, while one neuron with an initially discontinuous *f-I*-curve increased its onset firing frequency (Fig. 1D). Moreover, for a range of intermediate input currents, we observed episodes of spiking mingled with sub-threshold oscillations when GABA was added (Fig. 1B, orange-shaded region in Fig. 1C). Such stuttering behavior is indicative of noise induced transitions between coexisting bistable resting and spiking states (Izhikevich, 2010). GABA induced stuttering in *n* = 5 out of 7 neurons (Supplementary Fig. 1A).

**Figure 1:**
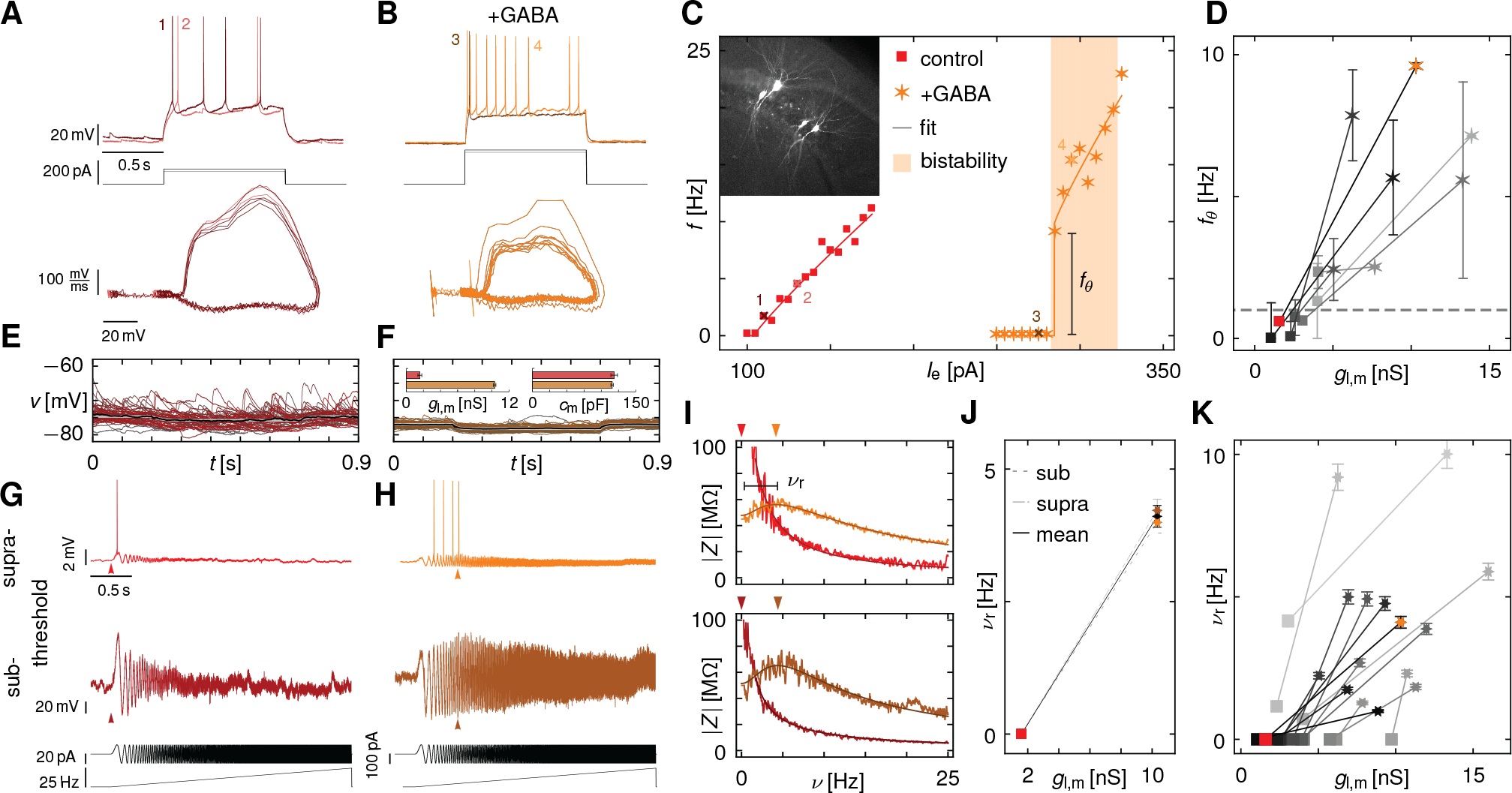
*Tonic GABA regulates firing rate encoding and resonance in CA3 hippocampal pyramidal neurons*. **(A, B)** Voltage traces (top) and phase portraits (*V*, 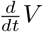-plane, bottom) of the AP dynamics in a CA3 neuron without (red, A) and with (orange, B) puff application of GABA in response to step currents. The AP onset firing rate increased with GABA and caused the neuron to stutter indicative of bistable dynamics. GABA also induced AP amplitudes to strongly decay during a spike train. **(C)** *f-I*-curve (dots) and region of stuttering/bistable dynamics (orange shading, 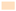) of the neuron in (A,B) without (red) and with GABA (orange). Fits to the *f-I*-curve (solid lines) show that GABA induced a non-zero onset spiking frequency *f*_*θ*_, switching the neuron from having a continuous to discontinuous *f-I*-curve. Inset shows stained CA3 neurons from recordings. **(D)** Onset firing frequency *f*_*θ*_ vs. measured resting state conductance *ɡ*_l,m_ across *n* = 7 CA3 neurons without (squares) and with GABA application (stars). Data from (A-C) in color. **(E, F)** Individual (red, orange) and mean (black) voltage responses of the CA3 neuron in (A,B) to *n* = 50 small hyper-polarizing step currents at resting potential without (E, red) and with (F, orange) GABA application. Input current induced depolarizations as well as amplitudes of spontaneous excitatory post-synaptic potentials decreased with little change in the resting potential. Fitting exponentials to the mean traces (gray) revealed an increase in the measured membrane conductance *ɡ*_l,m_ and no significant change in capacitance *c*_m_ due to GABA (insets). **(G,H)** Voltage responses (top, middle) of the CA3 pyramidal cell in (A) to a ZAP current with linearly increasing oscillation frequency from 0 to 25Hz (bottom). Responses just below (middle) and above AP threshold (top) for control conditions (G, red) and after GABA application (H, orange). **(I)** Impedance curves |*Z*| for the traces in (G,H) with fits to an RCL circuit (darker line) without (red) and with GABA (orange) just below (bottom) and above AP threshold (top). Arrows indicate resonance frequencies *ν*_r_ obtained from the fit (also shown in G,H). The response switches from an integrative to a resonant profile. **(J)** Peri-threshold resonance frequency *ν*_r_ obtained as the average of the resonance frequencies just below and above threshold as a function of *ɡ*_l,m_ for the neuron in (A) without (squares) and with GABA (stars). **(K)** Resonance frequency *ν*_r_ and leak conductance *ɡ*_l,m_ of all cells (*n* = 15). All non-resonant cells (*n* = 12) became resonant near theta frequencies, while intrinsic resonant cells (*n* = 3) increased their resonance frequency. Data from (D-J) in color.

To assess the effect of GABA on the biophysical properties of the pyramidal cells, we injected small hyper-polarizing step currents at resting potential to estimate the leak conductance *ɡ*_l,m_ and capacitance *c*_m_. For the neuron shown in Fig. 1A,B, application of GABA did not significantly change the resting potential or the capacitance, but increased the resting conductance by over four-fold and thereby reduced the amplitude of spontaneous postsynaptic potentials (Fig. 1E,F). Hence, the primary effect of GABA application is to introduce a shunt into the cell membrane. In all CA3 neurons GABA increased the leak conductance by a similar amount (*ɡ*_l,GABA_ = 5.9 ± 3.0nS). The increase in leak conductance *ɡ*_l_ correlated with the increase in AP onset frequency *f*_*θ*_ (*r* = 0.81, *p* < 10^−6^) (Fig. 1D).

To study how tonic GABA affects the resonance properties of CA3 cells near the AP generation threshold, we measured the frequency-resolved impedance *Z* (*ν*) by injecting sinusoidal currents whose frequency increased continuously in time (ZAP currents) (Hutcheon & Yarom, 2000). Before application of GABA, the voltage responses near but below AP threshold decreased as the oscillation frequency increased (Fig. 1G). Above threshold, a single AP was generated on the first peak in the stimulus. In both cases, the impedance curve decayed monotonically and had no detectable corner frequency (*ν*_r_ = 0Hz, Fig. 1I). As |*Z* (*ν*)| is inversely proportional to *ν*, the underlying electrical circuit is an integrator.

GABA changed the impedance profile and created a resonance (Fig. 1H), both below (*ν*_r_ = 4.0Hz) and above threshold (*ν*_r_ = 4.2Hz) (Fig. 1I,J). APs appeared near the resonance frequency (Fig. 1H). GABA induced resonances (*ν*_r_ = 4 - 10 Hz) in all previously non-resonant cells (*n* = 12) and increased *ν*_0_ in already intrinsic resonant cells (*n* = 3, Fig 1J). The strength of the resonances, as measured by the quality factor (*Q*-factor), varied from 1 to 2 (Supplementary Fig. S1B). The GABA-induced change in the resting membrane conductance was correlated with an increase in the resonance frequency (*r* = 0.59, *p* < 10^−4^) (Fig. 1K).

In summary, GABA generated an effective shunt in regular firing CA3 pyramidal cells, induced a discontinuity in the *f-I*-curve, decaying AP amplitudes, regions of bistable dynamics as well as peri-threshold resonances.

### Conductance clamp mimics the effects of GABA on AP dynamics and resonance

As GABA application increased the leak conductance in the CA3 cells (Fig. 1E,F), we hypothesized that the effects of tonic GABA application could be replicated by artificially changing the intrinsic membrane conductance by itself. We, therefore, added or subtracted a pure leak current

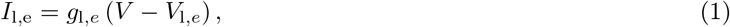

with membrane potential *V*, external leak conductance *ɡ*_l,e_ and leak reversal potential *V*_l,e_ to the cells using the dynamic clamp technique (Sharp et al., 1993) (Supplementary Fig. S2A,B). *V*_l,e_ was set to the measured resting potential of each cell.

Figure 2 shows that adding a leak current has the same effect as applying GABA. In all neurons (*n* = 18), the threshold for AP generation shifted with increasing leak, and neurons with initially continuous *f-I*-curve (*n* = 12) changed and exhibited a jump in the AP frequency at onset (Fig. 2A,B). Increasing *ɡ*_l,e_ also induced stuttering or bistable behavior over increasingly larger current ranges in these neurons (Fig. 2A,C). Both effects mimicked the effects of GABA application (Fig. 1A-C).

**Figure 2:**
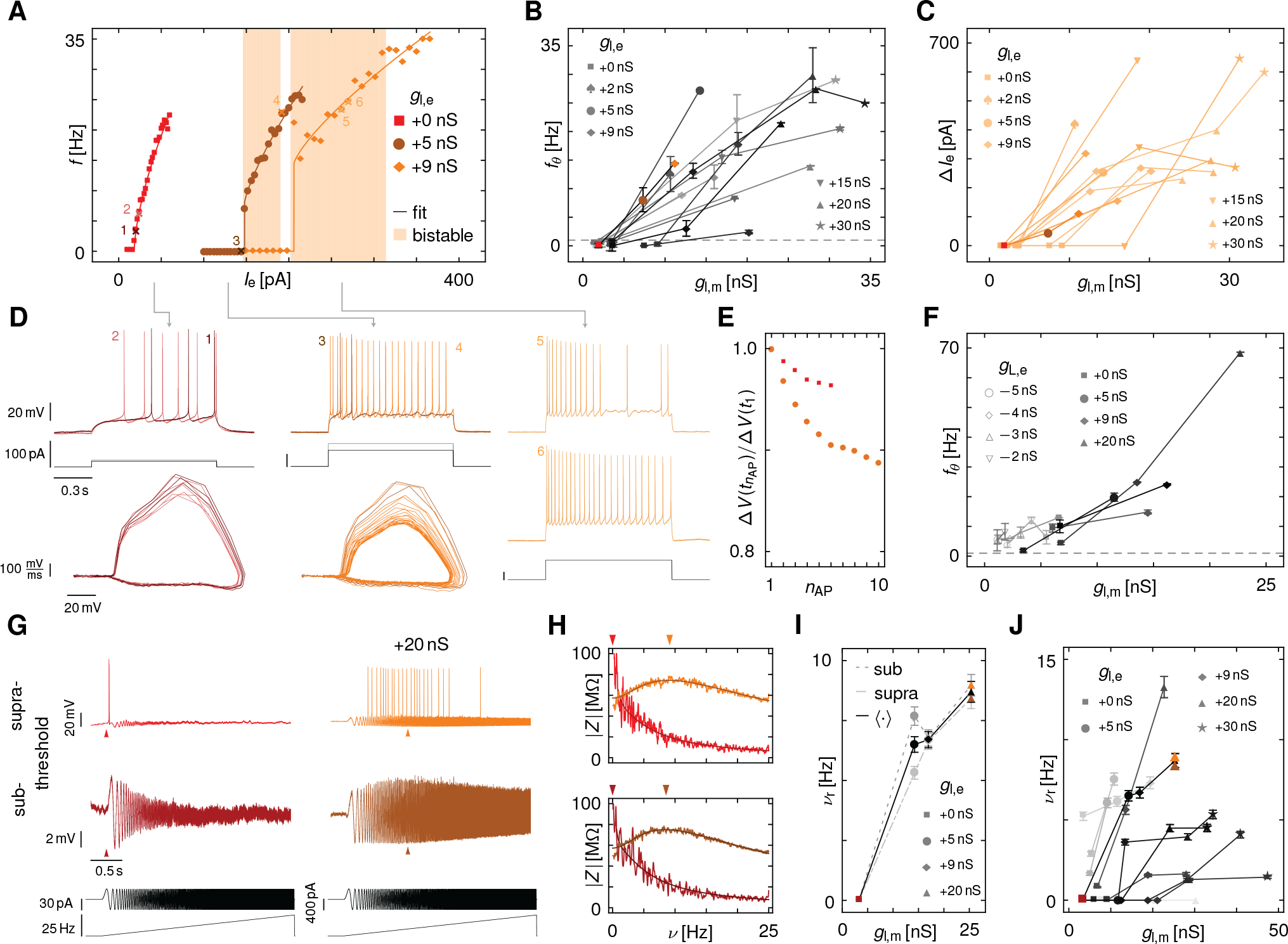
*Leak-conductance modulates firing rate encoding and resonance in CA3 hippocampal pyramidal neurons mimicking the effects of GABA*. **(A)** *f-I*-curves (dots) of a CA3 pyramidal neuron for different amounts of external leak conductance *ɡ*_l,e_ added in conductance clamp. Increased leak shifted the threshold for APs and induced stuttering or bistable dynamics (orange shading, 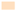). A fit to the *f-I*-curves (lines) shows a switch from zero to non-zero AP onset frequency *f*_*θ*_. Numbered crosses indicate the traces highlighted in (D). **(B)** Increase in onset firing frequency *f*_*θ*_ with increasing externally applied *ɡ*_l,e_ and measured leak-conductance *ɡ*_l,m_ at rest across different class-I neurons (*n* = 12). Data from (*A*) in color. **(C)** Width Δ*I*_e_ of the region of bistability for the cells in (B) systematically increased with external leak *ɡ*_l,e_. Data from (*A*) in color. **(D)** Voltage traces (top) and phase portraits (bottom) show the dynamics in response to step-currents (middle) for *ɡ*_l,e_ = 0 (left, red) and 5nS (middle, orange) of the neuron in (A). For *ɡ*_l,e_ = 9nS (right, light orange) two responses (top, middle) to the same step current injection (bottom) show periodic firing (top) or stuttering between spiking and peri-threshold oscillations (middle) in the region of bistability as indicated in (A). **(E)** AP voltage amplitudes vs. AP number *n*_AP_ in response to a step-current for *ɡ*_l,e_ = 0 (red) and 9nS (orange). **(F)** Onset spiking frequency *f*_*θ*_ for intrinsically class-II CA3 pyramidal neurons (*n* = 6) for different values of externally added and subtracted leak. For strongly negative external leak conductances, neuronal dynamics became unstable. **(G)** Voltage responses of a CA3 pyramidal cell to a ZAP current with linearly increasing frequency from 0 to 25 Hz (bottom) in 30 s. Responses just below (middle) and above spiking threshold (top) for *ɡ*_l,e_ = 0 (left, red) and 20nS (right, orange). **(H)** Impedance |*Z* (*ν*)| (line) for the traces in (G) with fit to an RCL circuit (darker line). Arrows indicate resonance frequencies *ν*_r_ of the fit. **(I)** Average *ν*_r_ of sub-and supra threshold resonance frequencies as a function of *ɡ*_l,e_ and *ɡ*_l,m_for all cells (*n* = 10), six of which did not intrinsically resonate. Cell in (G,H) highlighted in color. The resonances increased systematically with increasing leak.

Conductance clamp also caused similar changes in the voltage and phase-plane traces as GABA (Fig. 2D): during a train of action potentials, the voltage amplitude of the AP changed, with the first AP having the greatest height. Adding leak caused the height of APs to decrease more during the course of the spike train, while the approach to the “steady-state” AP height slowed down (Fig. 2E). With increasing leak conductance, the time from current onset to the first AP in these neurons strongly decreased (Supplementary Fig. S2C).

A smaller number of neurons intrinsically exhibited a discontinuous *f-I*-curve (*n* = 6, *f*_*θ*_ > 1 Hz at *ɡ*_l,e_ = 0 nS, Fig. 2F) and stuttering behavior. Addition of leak further increased the width of the bistability (Supplementary Fig. S2D). Subtracting leak reduced the onset firing frequency in these neurons (Fig. 2F), but did not induce a clear switch to a continuous *f-I*-curve. Decreasing the leak conductance even further finally destabilized the voltage dynamics, as the total leak conductance became zero or even negative.

Non-resonant neurons became resonant with increased intrinsic conductance, as revealed by injecting ZAP currents (Fig. 2G,H). This resonance could be measured both below and above spiking threshold (Fig. 2G-I). Induced or increased resonance frequencies were observed in all neurons measured (*n* = 10, Fig. 2I,J). The resonance frequency, measured at a baseline current of 95% of the threshold, increased significantly with the leak conductance (Supplementary Fig. 2E), while the strength of the resonance first increased and then decreased (Supplementary Fig. 2F,G). Interestingly, close to the leak-induced transition in the *f-I*-curve, the resonance appeared at sub-threshold voltages, and disappeared again close to threshold (Supplementary Fig. 2H,I).

In summary, adding leak conductance to CA3 cells via conductance clamp induced transitions in the *f-I*-curve’s excitability class, towards bistability and from integration to resonance mimicking the transitions induced via GABA. To understand these leak-induced changes in the AP dynamics in more detail we turned to study model neurons.

### Model neurons capture the effects of GABA shunt on AP dynamics and resonance

We fitted a standard CA3 pyramidal cell model (Migliore et al., 2010) to our recordings (cf. Methods). The resulting model recapitulates the main three experimental observations (Fig. 3): First, a region of bistability emerges as the GABA shunt increases, so that the addition of small noise fluctuations drives the neuron to alternate between periodic spiking and sub-threshold oscillations (Fig. 3A-C). Second, the excitability changes, with initially continuous *f-I*-curves giving way to discontinuous ones (Fig. 3A,B). Within a train of APs, constant-height APs turn into APs with successively smaller amplitudes (Fig. 3D). Third, the neuron’s transfer impedance switches from being low-pass to becoming resonant at a non-zero frequency (Fig. 3E-G, Supplementary Fig. 3A). The resonance frequency *ν*_r_ is a function of the holding membrane-potential *V*_hold_ (Supplementary Fig. 3B). In particular, for intermediate membrane shunts, the resonance frequency first rises and then falls as V_hold_ increases, similar to the experimentally found non-monotonic dependencies (cf. Supplementary Fig. S2H,I).

To understand the relationship between neuronal excitability, resonance, and bistability, we cast our computational CA3 pyramidal cell model into the general framework of conductance-based neuron models of the form

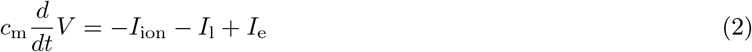

with *V* the membrane potential, *c_m_* the neuron’s capacitance, *I*_ion_ the trans-membrane currents of active ion channels, *I*_l_ the leak current, and *I*_e_ representing synaptic and externally applied currents. These types of models are amenable to a systematic analysis: Qualitative changes in the dynamics, such as the transition from resting to regular AP generation at a critical input current parameter *I*_e,*θ*_, are detectable as bifurcations. Generally, bifurcations separate different classes of dynamics in parameter space. Building upon an extensive body of earlier work (Rinzel & Ermentrout, 1989; Ermentrout, 1996; Izhikevich, 2010; Prescott et al., 2008a; Ermentrout & Terman, 2010), we performed a full bifurcation analysis on the fitted model (2) (cf. Methods), which revealed a complex sequence of dynamical transitions as the leak conductance was increased (Fig. 4).

**Figure 3:**
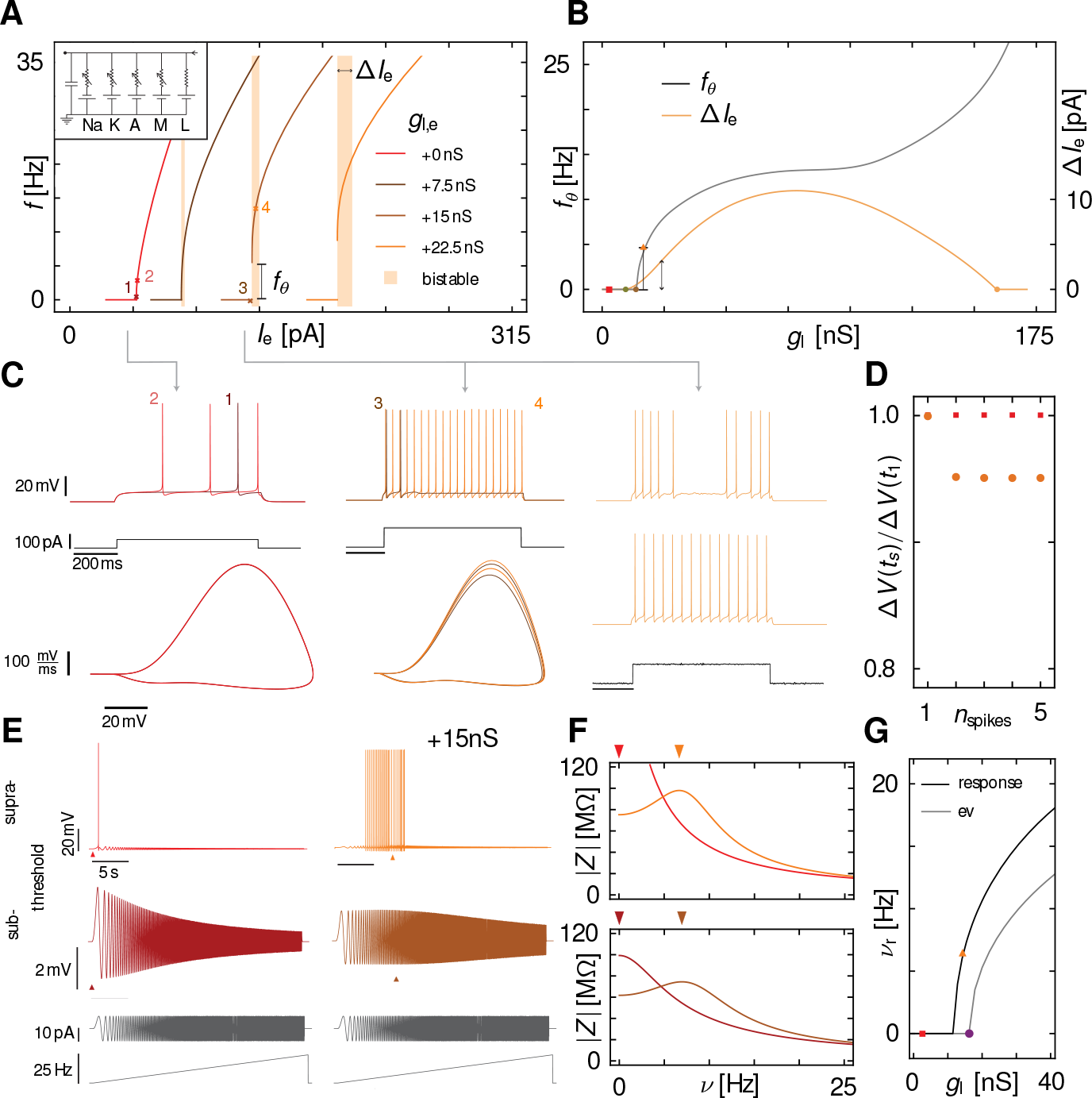
*Leak-induced transitions in AP dynamics and resonance in a CA3 model neuron*. **(A)** *f-I*-curves of a CA3 neuron model fitted to the experimental data (inset) for different external leak-conductances *ɡ*_l,e_. For increasing leak the AP onset frequency *f*_*θ*_ increases from *f*_*θ*_ = 0Hz to a non-zero value, and regions of bistability (orange shading, 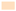) between quiescence and spiking emerge, similar to the effects of GABA and leak in the CA3 neurons (cf. Figs.1A,2A). **(B)** AP onset frequency *f*_*θ*_ at threshold and width Δ*I*_e_ of the bistable region as a function of the intrinsic leak *ɡ*_l_. Scale bars from (*A*). The neuron switches from class-I to class-II excitability 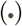 and from an integrator to a resonator 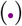 with increasing leak as observed experimentally (cf. Fig. 2B,C). Red square and orange triangle are the values for the traces plotted in **(C)**. Colored dots indicate co-dimension two bifurcations that organize the individual transitions (see text and Fig. 4). (C) Voltage traces (top) and phase portraits (*V*, 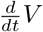-plane, bottom) of the AP dynamics in response to step currents (middle) without (*ɡ*_l,e_ = 0nS, red) and with (*ɡ*_l,e_= 20 nS, orange) added external leak which switched the neuronal excitability from class-I to II in (A). Numbers relate the traces to the values in (A). The traces on the right are the responses to a deterministic step current (top) and a step current with added noise (middle, bottom) for parameters in a bistable region. The noise induces switching between spiking and oscillatory fluctuations around the stable fixed point similar to the stuttering dynamics observed experimentally (Fig. 1D). **(D)** AP amplitudes for the traces in (C) are stable for the class-I model (red) and decay when adding leak (*ɡ*_l,e_ = 20nS, class-II, orange). **(E)** Supra-(top) and sub-threshold (middle) voltage responses to ZAP stimuli (bottom) for control (left) and with additional leak show a shift of the maximal response amplitude from 0 Hz (red arrow) to a non-zero frequency (orange arrow). **(F)** Impedance |*Z*| obtained from the responses in (E) change from input integration (red) to resonance (orange) with a non-zero resonance frequency *ν*_r_ (orange arrow) as in (B). **(G)** Resonance frequency *ν*_r_ (black) (from linear response at 95% of the AP onset current) and imaginary part of the eigenvalues of the stability matrix at AP onset (gray) both increase with leak in accordance with the experimental findings (cf. Figs. 1K,2J). Red square and orange triangle represent parameters from (E). The purple dot represents the parameter value of a Bogdanov-Takens bifurcation (cf. Fig.4).

**Figure 4:**
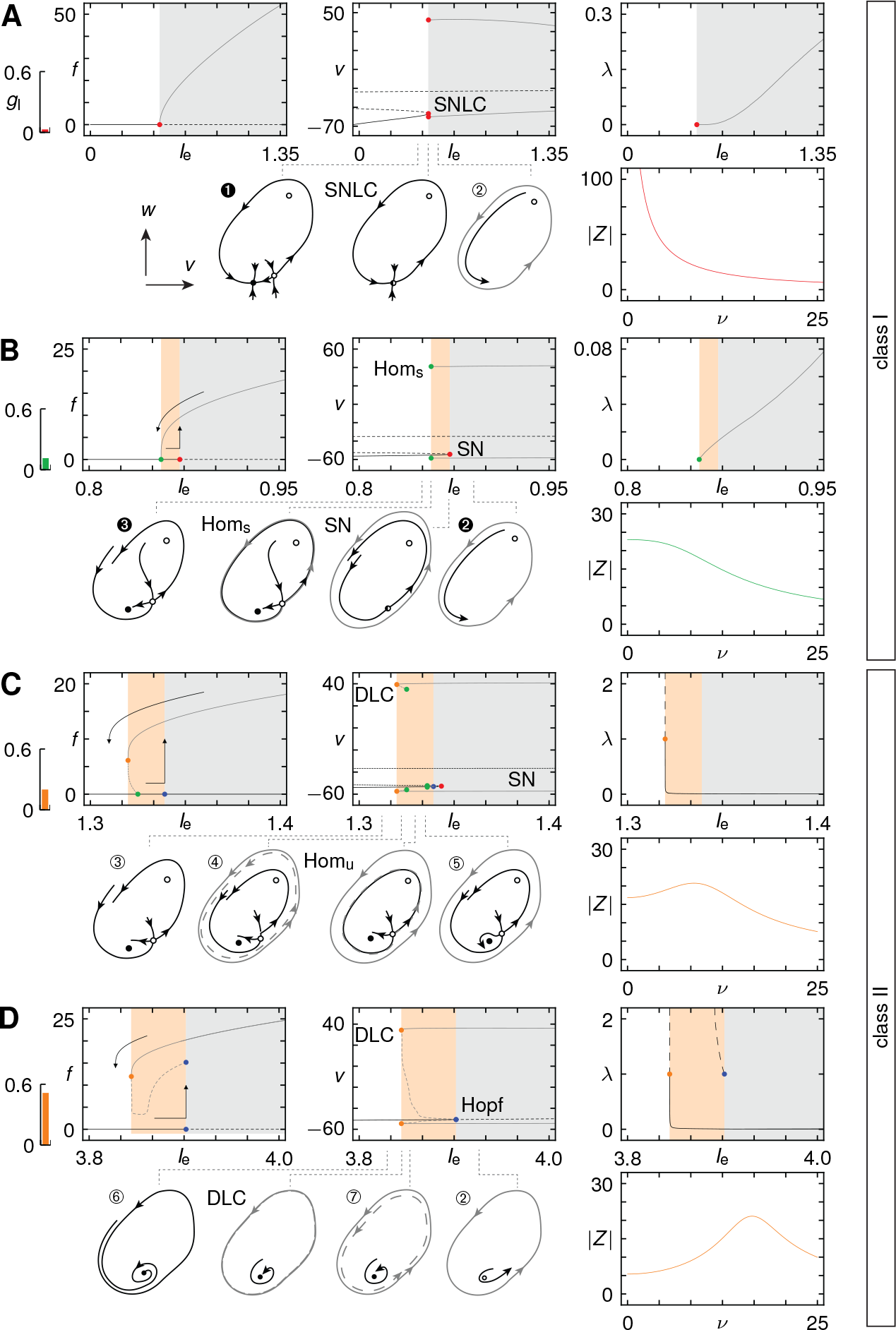
*A change in the dynamical transitions to regular AP generation underlie leak induced bistability, resonance and excitability transitions in a CA3 pyramidal cell model*. Each panel shows the *f-I-*curve (top left), the voltage *V* against the input current *I*_e_ (top middle), the impedance |*Z*| at 95% of the current at AP threshold (top right), sketches of the AP dynamics in a two-dimensional phase-plane of *V* and an effective activation variable *w* (bottom middle), and the largest non-trivial Floquet multiplier (FM) λ (bottom right) for the CA3 neuron model in Fig. 3. In the bifurcation diagrams, black lines - (––) indicate (un)stable fixed points, gray shading 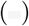 stable periodic AP generation, orange shading 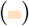 bistability in which a steady state and regular AP generation coexist. Gray lines (-) denote the maximum and minimum voltages of the stable AP cycle, dashing (––) indicates unstable orbits. In the sketches, closed (open) dots show stable (unstable) fixed points, half-filled dots saddle fixed points, black lines stable and unstable manifolds, gray lines periodic orbits. Units for the frequency and voltage are in Hz and mV. Units for the leak conductance and currents are in mS/cm^2^and *μ*A/cm^2^ or in 100 nS and 100pA for the neuron in Fig. 3 with a surface area *a* = 0.0001 cm^2^. **(A)** For a small intrinsic leak conductance *ɡ*_l_, a saddle-node on limit cycle bifurcation (SNLC, 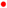) generates a continuous *f-I-*curve, the hallmark of class-I excitability. |*Z*| decays steadily, so there is no resonance. A small FM reflects fast attraction towards the limit cycle. **(B)** For intermediate *ɡ*_l_, the neuron shows a mixture of class-I and II excitability. A stable limit cycle arises through a homoclinic bifurcation (Horn, 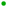); the resting state is destabilized for higher *I*_e_ in a saddle-node bifurcation (SN, 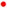). The resulting hysteresis in firing rates shows non-zero spike-frequency onset but a gradual decay to zero at offset. There is no resonance; the FM is zero at the bifurcation **(C)** For even larger *ɡ*_l_, class-II dynamics appear. A double limit cycle bifurcation (DLC, 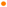) generates AP cycles with finite period, resulting in a discontinuous *f-I*-curve. The unstable cycle vanishes in a homoclinic bifurcation 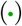, destabilization of the resting state is through a saddle-node, as before. The impedance |*Z*| shows a resonance close to threshold. At the bifurcation, the FM is unity, so attraction to the AP limit cycle is slow. **(D)** For large *ɡ*_l_, a sub-critical Hopf bifurcation (Hopf, 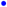) is observed together with a double limit cycle bifurcation (DLC, 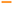), generating bistability, a more pronounced peri-threshold resonance, and discontinous class-II *f-I*-curves.

Under control conditions, the bifurcation from resting to regular AP generation in the CA3 model is governed by a saddle-node on invariant cycle bifurcation (Fig. 4A): In this scenario, a stable node corresponding to the resting state of the neuron merges with a saddle fixed point (Fig. 4A), so that both vanish. This results in sustained, repetitive APs. After the bifurcation, the dynamics remain slow in the vicinity of the disappearing fixed points, enabling the neuron to generate arbitrarily slow AP frequencies and the firing rate *f* increases continuously as 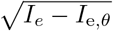 (Rinzel & Ermentrout, 1989; Ermentrout, 1996; Izhikevich, 2010; Ermentrout & Terman, 2010).

Hodgkin (Hodgkin, 1948) defined different classes of neuronal excitability: if the onset of action potentials is gradual in response to current injection, the cell belongs to class-I; if it is sudden and switch-like in frequency, the cell is in class-II; class-III comprises all cells that only fire a single AP or just a few APs in response to current injection. Based on this classification, the control CA3 neuron model exhibits class-I excitability.

The model is also non-resonant under control conditions. Indeed, if one measures the membrane-potential response to sinusoidal current stimuli with frequency *ν*, the complex-valued impedance *Z* (*ν*) behaves as 1/*i*ν** at low frequencies (Fig. 4A). As integration is mathematically equivalent to multiplication by 1/*i*ν** in frequency space, the neuron acts as an integrator of its synaptic inputs. In the absence of a GABA shunt, the experimental measurements of *Z*(*ν*) (Figs. 1G-K, 2G-J) are consistent with an integrator model.

Under constant current injection, the AP dynamics converges to a limit cycle (the steady-state AP waveform) at a rate measured by the Floquet multiplier λ. If λ is zero, then the dynamics approaches the limit cycle within one period of the oscillation; on the other hand, if λ is one, then there is no attraction. For class-I spiking, λ is close to zero at the threshold (Fig. 4A), which implies that the first AP already closes in onto the limit cycle immediately, so that subsequent APs in a spike train will have almost the same height and shape as the first AP, as observed experimentally in CA3 cells before GABA or a leak conductance was applied (Figs. 1A, 2C,D).

Increasing the leak conductance *ɡ*_l_ changes the bifurcations that give rise to regular AP firing (Fig. 4A-D). For intermediate *ɡ*_l_, the neuron shows a mixture of class-I and II excitability (Fig.4B): A stable limit cycle arises through a homoclinic bifurcation, while the resting state is destabilized at a higher input current *I*_e_ in a saddle-node bifurcation. Together, these two points give rise to a bistable region with a hysteresis in the *f-I*-curve: as *I*_e_ increases, the firing frequency jumps to a non-zero *f*_*θ*_ but as *I*_e_ decreases from above, the *f-I*-curve falls continuously to zero, scaling logarithmically near the homoclinic bifurcation (cf. Brown et al. (2004)). The impedance remains finite at zero frequency.

In the experiments, puff application of GABA added around 5 - 10 nS to the membrane conductance of CA3 neurons. Adding and equivalent amount to the model neuron again changes the dynamics (Fig. 4C): The neuron now exhibits class-II excitability both in the on- and off-set of AP generation. Stable AP firing appears via the simultaneous generation of a stable and unstable limit cycle in a double limit cycle bifurcation. The resting state is now destabilized via a Hopf bifurcation in which small-amplitude oscillations no longer decay but become unstable and result in APs. This Hopf transition also entails resonances manifested in a peaked impedance curve *Z*(**ν**) near the AP threshold. The rate of attraction to the stable limit cycle is slow, as measured by a non-zero Floquet multiplier. For even larger leak conductances, intermediate homoclinic bifurcations involving unstable limit cycles disappear and resonance frequencies increase (Fig. 4D).

A more complete picture of these transitions is obtained in a two-dimensional bifurcation diagram in (*I*_e_, *ɡ*_l_)- parameter space of the CA3 pyramidal neuron model (Fig. 5). In this diagram, the bifurcation points observed in Fig. 4 when varying the single parameter *I*_e_, now become lines separating different regions of qualitatively different dynamics (cf. Supplementary Fig. S5 for a full characterization of the dynamical regimes and sketches of the phase-plane dynamics). These bifurcation lines merge at higher-order bifurcation points: to reach these points, one needs to tune both *I*_e_ and *ɡ*_l_ and therefore are said to have co-dimension two.

**Figure 5:**
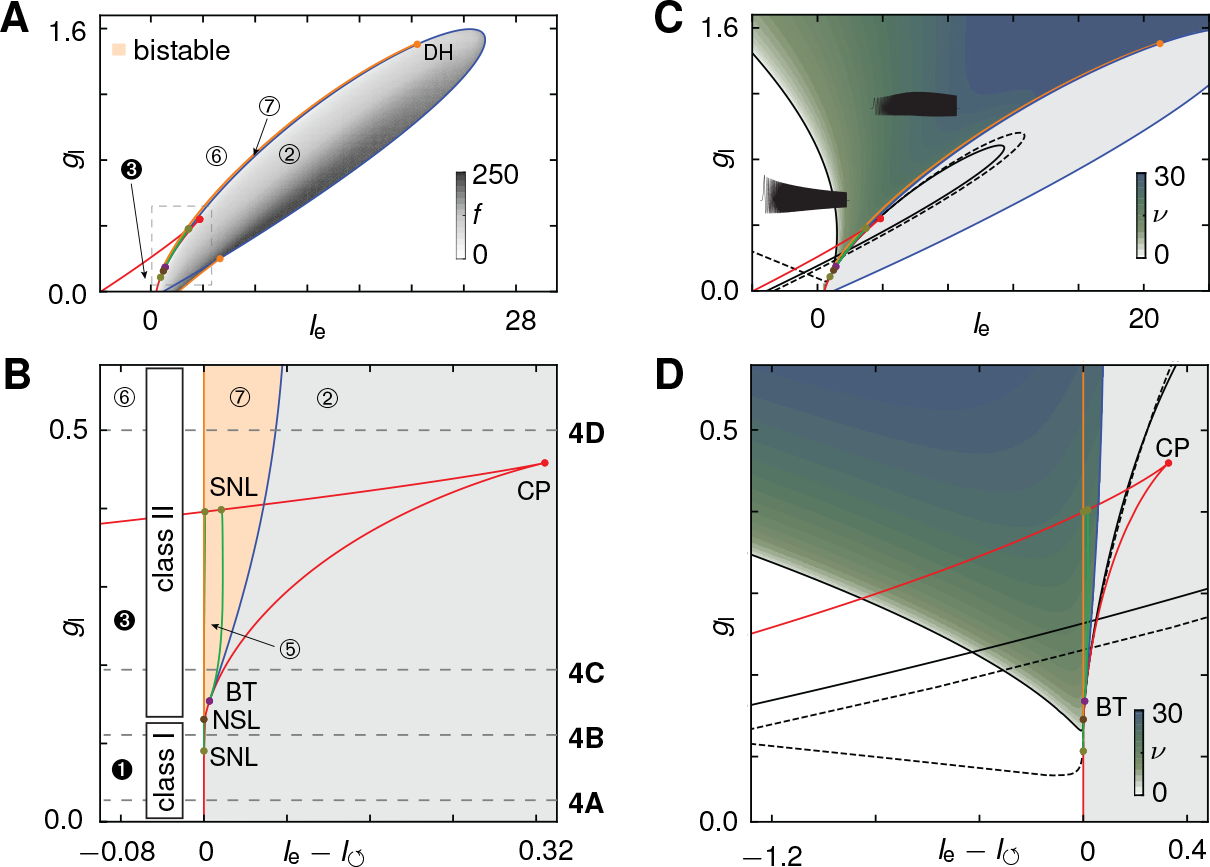
*Organization of leak-induced, transitions in neuronal excitability, bistability and resonance*. (**A**) Bifurcation diagram for the neuron model in Fig. 3 as a function of input current *I*_e_ and leak *ɡ*_l_. Gray level 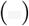 represents the frequency of periodic APs, orange shading 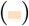 bi-stability between resting and spiking. **(B)** Fine structure of the transition resolved by shifting the input current *I*_e_ by the onset current for periodic AP generation *I*_↺_. Dashed horizontal lines refer to *ɡ*_l_ values used in Fig. 4. Class-I excitability due to a saddle-node on invariant cycle bifurcation (SNIC) bifurcation 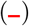, changes via the saddle-node-loop (SNL) point 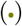 to a homoclinic bifurcation 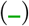; a neutral saddle loop (NSL) point 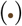 marks the transition to the double limit cycle (DC) bifurcation 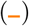 and class-II excitability. The region of bistability 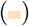 emerges at a SNL-point 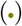 and vanishes at a degenerate Hopf (DH) point (only visible in A). At the Bogdanov-Takens (BT) point 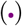, destabilization of the resting state changes from saddle-node 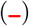 to Hopf 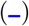 indicating a switch from integrative to resonant responses. **(C)** Leak-induced resonances appear near AP threshold and for increasing leak conductances extend into the sub-threshold regime. Colors and panels as in (A), green shading encodes resonance frequencies *ν*_r_. **(D)** Resonance behavior in the transition regime between class-I and class-II dynamics. At the black dashed line (––) the real eigenvalues of the linearized dynamics become imaginary, indicating onset of oscillatory components in the response. Black solid line (-) shows the transitions from zero to non-zero resonance frequency *ν*_r_. The transitions to resonance (-,––) pass tangentially through the BT-point (•). For *ɡ*_1_ below the BT-point and increasing *I*_e_, the resonance frequency *ν*_r_ first increases, then dips just before the AP threshold is reached. Units for the leak conductance and currents are in mS/cm^2^and *μ*A/cm^2^ or in 100 nS and 100 pA for the neuron in Fig. 3.

Enlarging the transition region (Fig. 5B) reveals that these co-dimension two bifurcation points mark the crucial changes in the dynamics: bistability emerges at a saddle-node-loop (SNL) bifurcation; class-I spiking changes to class-II at a neutral saddle loop (NSL) point; resonance at the threshold begins at the Bogdanov-Takens (BT) point. In the transition region from class-I to II, chimera-like states appear that have mixed properties, as two mechanisms for generating APs compete with each other: a pure threshold mechanism (saddle-node bifurcation) and an rapid amplification of peri-threshold oscillations (Hopf bifurcation). As the current *I*_e_ increases in this regime, the onset of APs can be class-II, while the offset of APs is class-I as the current *I*_e_ decreases.

The presence of an AP mechanism also affects the voltage dynamics below AP threshold. In the fitted CA3 model, resonances are observed in a wide swath of (*I*_e_,*ɡ*_l_)-parameter space (Fig. 5C,D). Interestingly, the line that separates the non-resonant from the resonant regime passes through the BT tangentially and thereby gives rise to a resonant region below it in which the resonance frequency *ν*_r_ first increases and then dips just before the AP threshold (Fig. 5D).

Detailed analysis of the CA3 model neuron thus revealed a sequence of bifurcations that control the complex dynamical transitions observed in response to application of GABA or anincrease in leak conductance. Varying tonic GABA, therefore, systematically controls the AP dynamics and resonance properties in CA3 neurons.

### A-type currents enable tonic GABA-mediated modulation of AP dynamics

Fitting the CA3 pyramidal cell model to the experimental traces required adjusting the A- and M-type potassium currents to activate close to AP threshold. So what is the role of these currents in shaping the transitions in the AP dynamics?

Removing the A-type conductance from the model leaves the overall structure of the bifurcation diagrams unchanged (Fig. 6A). However, the parameter values for the degenerate bifurcations points (SNL, NSL, BT), which organize the individual transitions, do shift towards higher leak conductances. This shift systematically increases with decreasing maximal A-type conductance *ɡ*_A_ (Fig. 6B). Notably, in the absence of the A-type current, the increase in leak conductance mediated by the application of GABA (cf. Fig. 1) is not capable of inducing the dynamical transitions in the model.

When we remove A- and M-type currents from the neuron model, we are left with only AP generating currents. Remarkably, the bifurcation structure for this model still remains unchanged but with an even larger shift towards higher leak conductances (Fig. 6C). In particular, sub-threshold resonances still exist, and thus, the specialized sub-threshold currents such as *I*_h_, *I*_M_, or *I*_Nap_, while often shaping resonances, are not necessary to induce these. A-type as well as M-type conductances, by activating close to AP threshold, effectively add to the overall leak and GABA shunt. This additional shunt effectively reduces the amount of leak needed to induce the transitions in excitabiltiy and resonance. Near the AP threshold, the A-type conductance contributes the largest amount of additional net conductance (Fig. 6D).

While M-type currents are known to contribute to higher leak conductances and support class-II excitability (Prescott et al., 2006), the A-type current is usually thought to delay APs, thereby reducing firing rates and moving class-II neurons towards class-I excitability (Connor & Stevens, 1971; Ermentrout, 1996). Here, surprisingly, by contributing a major component to the overall leak conductance at threshold the situation is reversed: The A-type current enables GABA to modulate AP dynamics from class-I to class-II, towards resonance and bistability.

### The modulation of AP dynamics follows a universal topology

Intrigued by the similarity of the bifurcation diagrams when varying A-type and M-type currents (Fig. 5,6) we sought to address the generality of our results. Indeed, a large number of non-bursting neuron models, ranging from fast-spiking inter-neurons (Wang & Buzsáki, 1996; Erisir et al., 1999) to slow muscle fibers (Morris & Lecar, 1981) all share the same topological structure of the bifurcation diagrams (Fig. 7A,B, see Supplementary Fig. S7i for more examples). What is the underlying reason for this prevalence of the transition structure?

**Figure 6:**
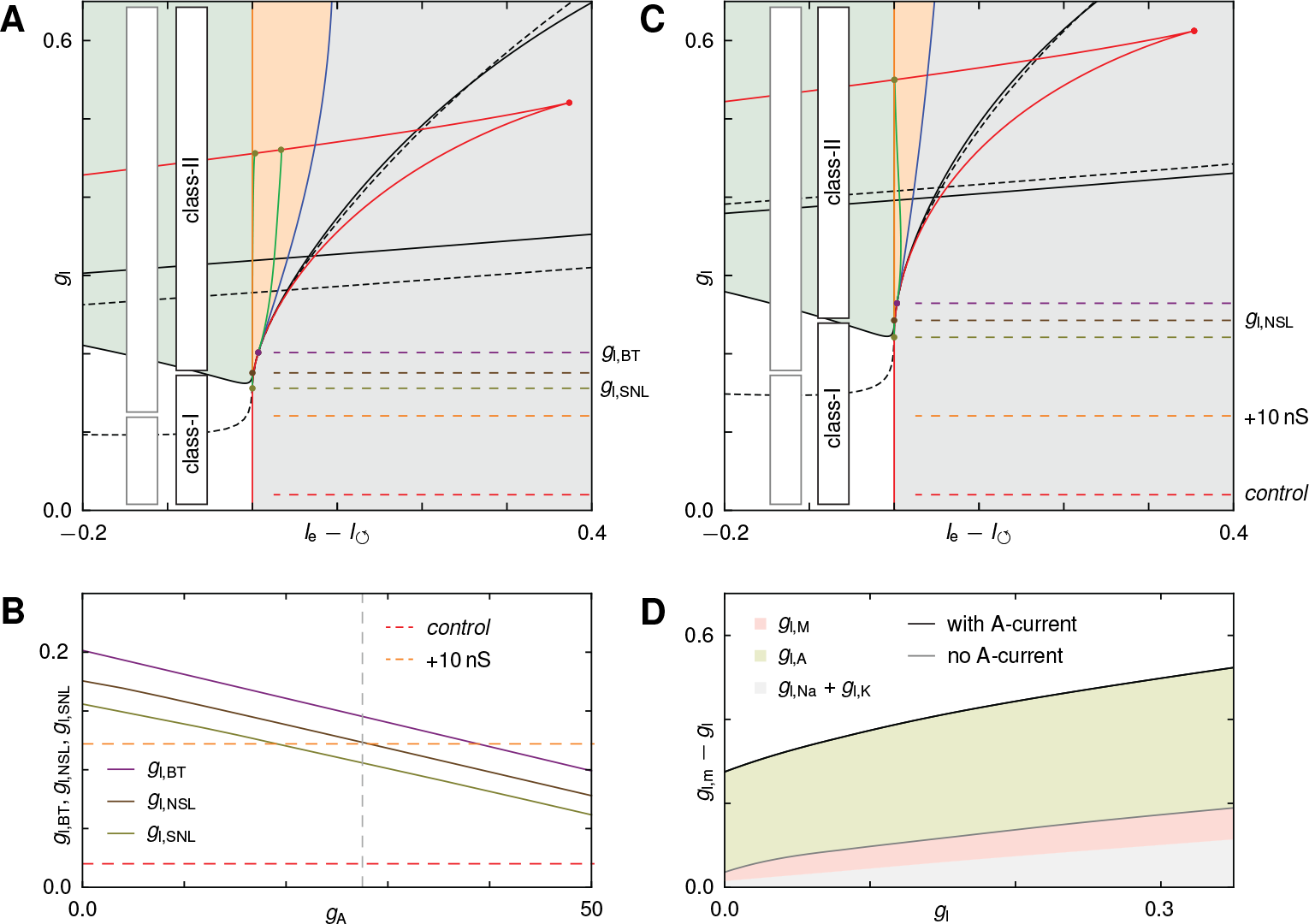
*Transient A- and M-type currents facilitate* GABA-*mediated modulation of AP dynamics*. **(A)** Bifurcation diagram for the CA3 neuron model in Fig. 3 without the A-type potassium current (*ɡ*_A_ = 0)- The topology of the diagram remains the same as in Fig. 5B, but a larger *ɡ*_l_ values are required to reach the critical points that organize the dynamical transitions to resonance (*ɡ*_l,BT_), bistability (*ɡ*_l,SNL_), and from class-I to II excitability (*ɡ*_l,NSL_ black boxes). For comparison the class-I to II regions in the full model from Fig. 3B are shown (gray boxes). Adding *ɡ*_l,e_ = +10 nS external leak conductance (orange dashed line) as caused e.g. by tonic GABA application (cf. Fig. 1) to the control conditions (red dashed line) no longer is sufficient cross the critical leak conductances to switch to resonance and class-II excitability (cf. Fig. 5). **(B)** Leak conductances for the bifurcation points as a function of the maximal conductance *ɡ*_A_ of the A-type current in the CA3 model neuron (cf. Fig. 3). Increasing *ɡ*_A_ reduces the critical thresholds for the individual transitions. Only for sufficiently strong A-currents (gray dashed line indicates *ɡ*_A_ for the model in Fig. 3) does an additional external leak of *ɡ*_l,e_ = 10 nS (orange dashed line) as observed e.g. by addition of GABA (cf. Fig. 1) change the conductance of sufficient magnitude to cause transitions in the dynamics. **(C)** Bifurcation diagram in the absence of both the A-type and M-type potassium currents (*ɡ*_A_ = *ɡ*_M_ = 0). The critical leak conductances of the transition points increase further including the transition between excitability classes (black boxes) as compared to (A) (gray boxes). **(D)** Measured total leak conductance *ɡ*_l,m_ in the CA3 model when the voltage is at 95% of the AP threshold plotted as a function of the intrinsic leak conductance *ɡ*_l_ with (black) and without (gray) A-current in the CA3 model of Fig. 3. The active ion channels are partly activated and thus effectively increase the measured leak conductance. The A-current provides the largest contribution to the total leak conductance (*ɡ*_l,A_, green shading) in comparison to the M-type (*ɡ*_l,M_, red shading) and AP-generating currents (*ɡ*_l,Na_ + *ɡ*_l,K_ shading). Units for the leak conductance and currents are in mS/cm^2^and *μ*A/cm^2^ or in in 100 nS and 100 pA for the neuron in Fig. 3.

**Figure 7:**
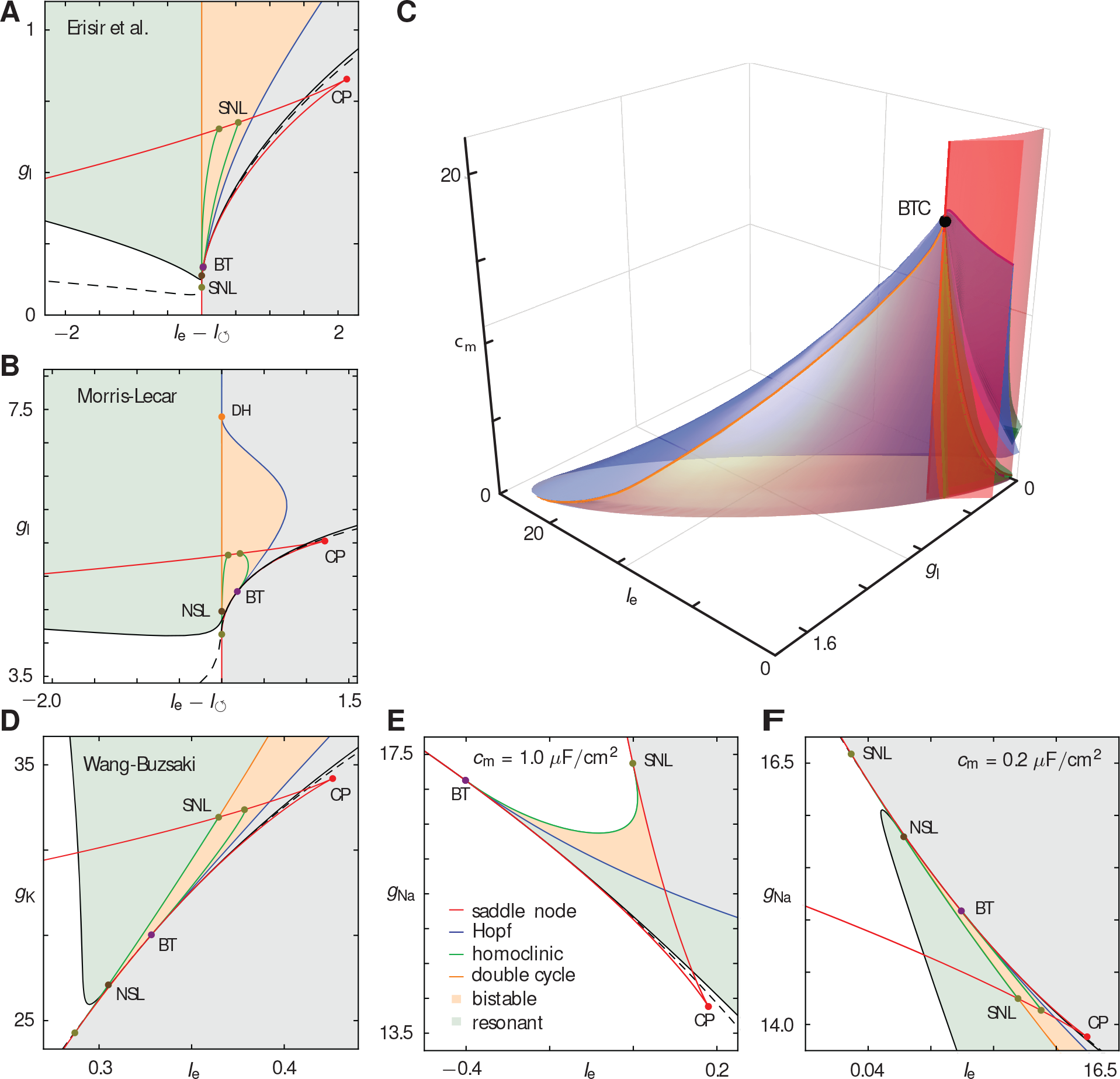
*A universal structure underlies the modulation of AP generation, bistability and resonance*. **(A,B)** Two dimensional bifurcation diagrams for a fast spiking inter-neuron model Erisir et al. (1999) (A) and a generic spike generator Morris & Lecar (1981) (B). Colors, lines and dots as in Fig. 6A. The diagrams have the same structure as the diagrams for the CA3 model in Figs. 5 and 6. **(C)** Three parameter bifurcation diagram for the CA3 neuron model in Figs. 3-6 for the input current *I*_e_, the leak conductance *ɡ*_l_, and the capacitance *c_m_*. The bifurcation lines become surfaces and the co-dimension two bifurcations becomes lines (colors as lines and points in Fig. 5). All lines and surfaces converge towards a degenerate co-dimension 3 Bogdanov-Takens-cusp bifurcation (BTC, •). **(D)** Bifurcation diagram for another generic neuron model Wang & Buzsáki (1996) varying the maximal potassium conductance *ɡ*_K_ together with the input current *I*_e_. The diagram shows the same topological transition structure as observed when varying the leak conductance as e.g. in (A,B). **(E)** Bifurcation diagram for the generic neuron model in (D) for the maximal sodium conductance *ɡ*_Na_ and the input current *I*_e_. The diagram differs in its structure. However, the structure can be identified in the full BTC structure as a horizontal section through the diagram in (C) when choosing the capacitance above the BTC point (cf. also Supplementary Fig.7iii). **(F)** Diagram as in (B) but with a fixed lower capacitance giving rise to a transition structure that is topologically the same as the generic structure identified before. Due to the opposing direction of the Sodium current in comparison to the Potassium current the diagram is inverted on in the *ɡ*_Na_ axis. (D-F) indicated that the BTC-point very generally organizes transitions in AP generation also when varying other bio-physical parameters. Units for the leak conductance, currents and capacitance are in mS/cm^2^, *μ*A/cm^2^, and *μ*F/cm^2^.

An answer can be found by including the membrane capacitance *c_m_* as a third generic parameter in the bifurcation analysis. We obtain three-dimensional (*I*_e_,*ɡ*_l_,*c*_m_)-parameter bifurcation diagrams in which the dynamical regimes are now separated by bifurcation-surfaces (Fig. 7C). The degenerate points in the two dimensional diagrams now correspond to lines at which the bifurcation surfaces intersect. By taking sections through these three-dimensional diagrams at fixed *c_m_*, we recover the two-dimensional diagrams for the shunt-induced transitions (cf. Fig. 5). Remarkably, a combination of parameter values exists for which the bifurcation surfaces and lines all collapses into a single degenerate bifurcation point (Fig. 7C): a Bogdanov-Takens-cusp (BTC) point (Dumortier et al., 1991). In general, three sub-types of BTC point exist (Dumortier et al., 1991), but we find that class-I excitable models either have a BTC point of focus or elliptic type (Supplementary Text and Table S1).

According to multiple-bifurcation theory (Guckenheimer, 1986), degenerate bifurcation points unfold in a generic way when the parameters are changed, thereby fully determining the bifurcation diagrams in its neighborhood. For example, an ordinary Bogdanov-Takens point entails the presence of saddle-node, Hopf and homoclinic bifurcations in its vicinity (see e.g. Fig. 7A). All bifurcation surfaces and lines observed in Fig. 7C are indeed predicted by the universal unfolding of the degenerate BTC point (Dumortier et al., 1991); hence, the BTC point acts as an organizing center (Guckenheimer & Holmes, 1983) for the complex transition structure observed across various neuron models.

Now consider a generic conductance-based neuron model of the form (2) with arbitrary active ion-channel conductances. Interestingly, we can mathematically prove (cf. Methods and Supplementary Text) that if such a model exhibits class-I excitability it will always have a BTC point in (*I*_e_,*ɡ*_l_,*c*_m_)-parameter space (Theorem 1). This BTC point will be of focus or elliptic sub-type, provided a generic condition (10) is fulfilled (Proposition 1). As a consequence, increasing the membrane conductance in class-I neurons will always force a transition to class-II behavior, induce bistability, and cause resonance. Further experiments supported this claim: when repeating our dynamic clamp experiments on a completely different set of regularly firing neurons, recorded not from the CA3 region of hippocampus but from the gerbil’s dorsal nucleus of the lateral lemniscus (DNLL), we observed the same transitions in the dynamics upon increasing the leak conductance (Supplementary Fig. S7ii).

Consistent with our experimental observations (Fig. 2F), the converse is not necessarily true: for some class-II neuron models, the transition to class-I behavior would occur at negative leak conductances, which destabilize the dynamics: For example, for the class-II Hodgkin-Huxley model (Hodgkin & Huxley, 1952), only a BTC point of saddle type exists, which entails regimes with unbounded dynamics (Dumortier et al., 1991) (Supplementary Table S1).

The results above generalize even further to transitions induced by other parameters in the conductance-based models: once the BTC-point is identified in (*I*_e_,*ɡ*_l_,*c*_m_)-space, it will unfold into any two parameter directions. The bifurcation diagrams will still be sections through a 3D diagram as in Fig. 7C. Indeed, the unfolding theorem correctly predicts the topology of the bifurcation diagrams when varying the peak conductances of the AP potassium and sodium current or their half-activation voltages (Fig. 7D-F, Supplementary Fig. S7,iii,iv). Interestingly, certain parameter diagrams give rise to an altered fine structure that corresponds to sections with fixed *c*_m_ above the BTC point (Fig. 7E, Supplementary Fig. S7iii). Nevertheless, these diagrams show the same sub-transitions from class-I to II excitability, from integration to resonance, as well as towards a region of bistability.

Taken together, these results provide strong evidence that AP dynamics are generically organized by a degenerate BTC-point. Neuro-modulators, by affecting the AP conductances, can, therefore, move the neuron systematically into the different dynamical regimes that exist around the BTC-structure. Near the BTC-point the normal form equation (10) describes these dynamics, regardless of the details of the underlying model (Dumortier et al., 1991). The actions of neuro-modulators map onto changes of the unfolding parameters in (10). Thus this equation may provide a first step towards a general theoretical framework for the neuro-modulation of APs and serve as a starting point to study network effects of AP-modulation.

**Figure 8:**
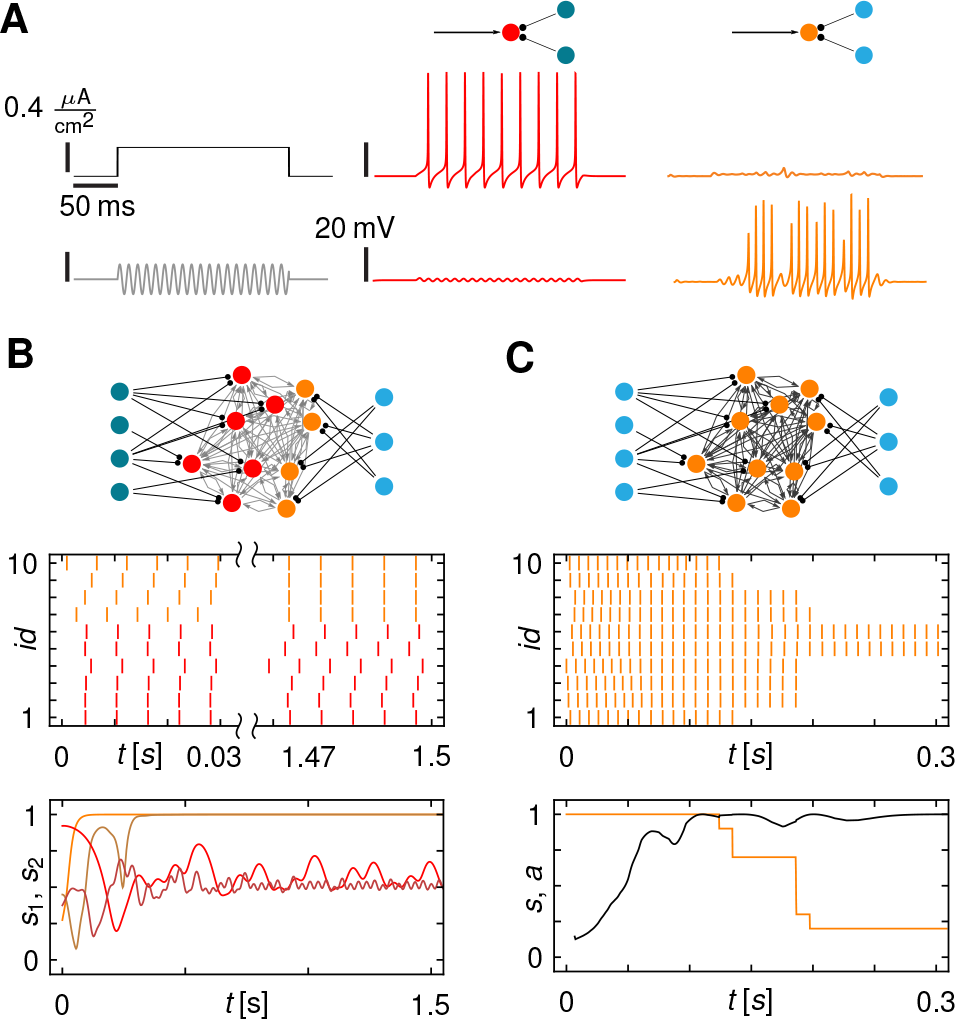
*Dynamic control of neuronal excitability and resonance enables flexible signal gating, dynamic grouping and limits epileptic like synchronization*. **(A)** A class-I excitable model neuron (red or orange disk) receiving an external input current (arrow) and synaptic inhibition from inter-neurons (dark or light turquoise disks). For inactive inhibitory neurons (middle column) the target neuron is non-resonant and spikes in response to a constant (middle panel) but not to oscillatory input (bottom). Activation of the inhibitory synapses (right column) at a Poisson rate of 40Hz induces resonance in the center neuron (orange disk). Stimulation with an oscillatory stimulus near the resonance frequency induces spiking. Fluctuations in the response are due to stochastic activation of the inhibitory neurons. **(B)** Network (top) of *N* =10 excitatory neurons (red) receiving synapses from two groups of inhibitory neurons. For silent inhibitory neurons (dark turquoise), the excitatory neurons are class-I (red) and desynchronize (Supplementary Fig. S8A). Activating a subset of inhibitory neurons (light turquoise) provides sufficient shunting to dynamically change a subset of postsynaptic neurons to class-II (orange), giving rise to a hybrid class-I/II network. The subset of class-II neurons, initially asynchronous, synchronize (orange) while the class-I neurons, initially synchronous, desynchronize (red). *s*_1_ (red line) and *s*_2_ (orange line) are synchrony measures of the individual sub-groups. Random initial conditions lead to the same dynamic grouping (bottom, dark red and orange lines). Switching all neurons to class-II by activation of all inhibitory neurons lead to network wide synchronization (Supplementary Fig. S7B). **(C)** Bistability controls the maximal number of synchronized neurons. Network as in (B) with increased coupling weights and all inhibitory neurons active. After an initial synchronization phase, the combined synaptic inputs are sufficiently strong to push neurons from spiking into the basin of attraction of the coexisting fixed point, leaving only a smaller fraction *a* of active neurons (orange line at bottom).

### Synaptic induced AP modulation flexibly controls signal gating, sub-ensemble formation, and limits on synchronization

How would synaptically controlled changes in the AP dynamics affect neuronal function? In Fig. 8A, spiking inhibitory neurons cause a postsynaptic neuron to resonate through slow shunting inhibition (as in Fig. 3), thereby regulating the postsynaptic neuron’s spiking response to constant and oscillatory stimuli. Controlled induction and tuning of resonances thus can flexibly gate oscillatory inputs.

As inhibitory networks are thus capable of modulating the neuronal excitability of their synaptic targets, they could act as a synchrony switch for interconnected principal neurons, given that class-I neurons tend to desynchronize, while class-II neurons synchronize (Ermentrout, 1996). We simulated networks of excitatory class-I neurons that were the targets of interneurons with slow shunting synapses. When the interneurons were silent, the excitatory neurons were class-I and desynchronized (Supplementary Fig. S7A). Inhibition effectively increased the leak conductance of the excitatory neurons and switched them to class-II, causing them to synchronize (Supplementary Fig. S7B,C). Interestingly, if only a subgroup of the excitatory neurons switched to class-II through inhibition, we observed that only the neurons in this sub-ensemble synchronized their spikes (Fig. 8B). The others became asynchronous despite the stronger synchronous input to all neurons.

More interestingly, pushing neurons into the bistable class-II regime induced synchronization but also limited the total number of active neurons (Fig. 8C): once the synchronized synaptic input pulses became strong enough, they drove spiking neurons into the basin of attraction of the co-existing resting state (Supplementary Fig. S7D). This process continued until the combined pulse strength of the active neurons became too weak to silence even more neurons. Together, synchronization and bistability effectively constrained the total number of simultaneously active neurons.

Modulating the AP dynamics of neurons via shunting inhibition thus provides a basic biophysical mechanism to control signal processing and collective network dynamics. In particular, this dynamic neuronal excitability can selectively gate oscillatory inputs and bind a self-limited number of neurons by synchrony (Hebb, 1949; von der Malsburg, 1991), which has potential roles in flexible neuronal coding and information transmission (Fries, 2005).

## Discussion

We have shown that activating GABA receptors caused CA3 pyramidal neurons to develop membrane-potential resonances, bistability, and shift their excitability from continuous class-I to discontinuous class-II *f-I*-curves (Fig. 1). This effect can be mimicked in conductance clamp, both in CA3 and in the brainstem (Figs.2, S7ii). Modeling revealed a sequence of specific bifurcations underlying the observed transitions (Fig. 3-5) and showed that the presence of transient potassium A-currents poised the model neuron at the transitions’ edge, so that the GABA shunt is capable of controlling neuronal excitability (Fig. 6). Theory showed that the sequence of dynamical transitions is universal (Fig. 3-7, Theorem 1) and can be invoked by changing any malleable biophysical property of the neuron (Fig. 7). These results may thus serve as starting point to develop a general theoretical framework for the modulation of AP dynamics and its role in neural computation (Fig. 8).

### Shunting through GABA modulates AP generation dynamics and induces resonances

GABA-ergic inhibition, in addition to controlling overall network activity and taming epileptic-like discharges (Semyanov et al., 2004; Mody & Pearce, 2004; Jeong & Gutkin, 2007; Olah et al., 2009; Deng et al., 2009; Brickley & Mody, 2012; Ammer et al., 2012; Pavlov et al., 2014), can also enhance network oscillations by alternating phasically with excitation (Cobb et al., 1995; Mody & Pearce, 2004; Vida et al., 2006; Mann & Paulsen, 2007; Jeong & Gutkin, 2007; Ammer et al., 2012; Stark et al., 2013; Pavlov et al., 2014). Here we demonstrated, that GABA also changes the postsynaptic neurons’ excitability class, induces membrane potential resonances, and thereby can reinforce network synchronization.

In CA3 pyramidal cells, GABA induced resonances in the range of 3-8 Hz, which overlaps with hippocampal theta-rhythms (Buzsaki, 2002). GABA, by controlling the leakiness of a neuron, determined the neuron’s resonance frequency, both above and below the AP threshold (Figs. 1, 5). The precise frequency also depends on the AP generator (Supplementary Fig. S7i): in contrast to the CA3 model, a model for fast-spiking interneurons (Wang & Buzsáki, 1996) predicts that the AP mechanism produces peri-threshold resonances in the gamma range of 30-200 Hz.

Sub-threshold resonances can arise through the interaction of specialized currents such as persistent sodium, low-threshold calcium, mixed-cation, and hyperpolarization-activated potassium currents (White et al., 1995; Hutcheon & Yarom, 2000; Hu et al., 2002; Stark et al., 2013; Rotstein & Nadim, 2014). Those resonances persist after blocking AP with TTX (Pike et al., 2000). Here we showed, however, that none of these specialized currents are necessary for resonance. In the CA3 model, the peri-threshold resonances remained after stripping away all non-AP-generating currents, including the M-current (Fig. 6C). Hence, the mere presence of an AP-generating mechanism proves to be sufficient for a membrane potential resonances. Instead of attenuating oscillations and resonance, here membrane shunt in combination with the AP generating conductances actually evinced them. As sub-threshold resonances generated by specialized currents impact the spiking statistics (Richardson et al., 2003; Broiche et al., 2012) resonances due to the AP generator likely play a similar important role in determining spike timing and information transmission (Llinás, 1988; Alonso & Llinas, 1989; Hutcheon & Yarom, 2000; Buzsaki, 2006; Haas & White, 2002; Vaidya & Johnston, 2013).

We treated the effect of GABA as a membrane shunt, ignoring its inhibitory or excitatory effects that occur in other circuits or at different developmental stages (Ben-Ari, 2002; Gulledge & Stuart, 2003; Pavlov et al., 2014). Yet these GABA currents can be accounted for by splitting *I*_l_ = *ɡ*_l_ (*V* − *V*_GABA_) into two components: a shunt *ɡ*_l_ (*V* − *V*_l_) with respect to the resting membrane potential *V*_l_ and a voltage-independent current *ɡ*_l_ (*V* − *V*_GABA_) that depends on the reversal potential *V*_GABA_ of the GABA-ergic conductances. The second term can be subsumed into the external current parameter *I*_e_, resulting in an (*I*_e_, *ɡ*_l_)-bifurcation diagram whose qualitative structure is unchanged: An increase in *V*_GABA_ only tilts it towards the left. Moreover, the same arguments apply to other passive-like conductances, e.g. due to Potassium leak channels (Goldstein & Bockenhauer, 2001) or background network activity (Destexhe & Paré, 1999; Prescott et al., 2008b).

GABA conductances have the highest density in the peri-somatic region (Megias et al., 2001) and extra-synaptic GABA currents can show an 8-fold increase in the presence of additional neuro-modulators (Wohlfarth et al., 2002), making GABA a particularly suitable neurotransmitter to regulate AP dynamics. Modeling predicts that also the neurotransmitter dopamine can control AP excitability (Morozova et al., 2016). Generally, controlling AP dynamics, via GABA, dopamine or other transmitters, thus provides a powerful mechanism to regulate neuronal coding.

### A-currents enable GABA-mediated transitions in AP dynamics and resonance

A- and M-type potassium currents facilitate dynamical transitions in CA3 pyramidal cells, as these currents activate near AP threshold and provide additional shunting. Indeed, without these currents amplifying the effect of GABA, the induced conductance is not necessarily of sufficient magnitude to cause transitions in the dynamics from class-I to II excitability and resonance (Fig. 6).

The role of A-type currents in supporting GABA-induced class-II excitability stands in contrast to it’s classical role: its slow inactivation delays spike onset, linearizes the *f-I*-curve, and ensures class-I excitability (Connor & Stevens, 1971; Connor et al., 1977; Ermentrout, 1996; Rinzel & Ermentrout, 1989; Rush & Rinzel, 1994). Intriguingly, large A-type conductances by themselves have been shown to induce class-II excitability (Drion et al., 2015), this time through a third mechanism organized by a pitchfork bifurcation (Franci et al., 2012, 2013).

Surprisingly, these seemingly different effects can be understood in terms of the BTC-point unfolded in the A-type conductance parameter *ɡ*_A_ and input current *I*_e_: An initial increase in *ɡ*_A_ follows the sequence of transitions from class-II to I as observed in Fig. 7F (sketch in Supplementary Fig. S5) for the *ɡ*_K_ conductance. Increasing *ɡ*_A_ further triggers a transition back to class-II excitability starting with a sequence as seen in Fig. 7E (sketch in Supplementary Fig. S7iii). Geometrically, the surface of the BTC unfolding in (*ɡ*_A_, *I*_e_)-space bends around the BTC-point in a ∪-shaped manner and thereby captures the opposing effects when changing the A-type conductance. This example highlights the utility of our approach to systematically understand AP modulation.

### Biophysical mechanisms of transitions in AP dynamics

In contrast to our analysis using multiple bifurcation theory, previous research has sought to explain the difference between the two neuronal excitability classes qualitatively, using the steady-state *I-V*-curve (Rinzel & Ermentrout, 1989; Izhikevich, 2010; Prescott et al., 2008a):

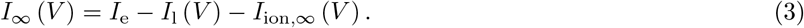

Here *I*_ion,∞_ (*V*) are the active ion currents at steady state. Class-I neurons usually have a kink in the steady-state *I-V*-curve (see also Supplementary Text, Lemma 5), whereas class-II neurons do not. The lack of a kink in class-II neurons implies that positive feedback must race against negative feedback to produce an AP; as a consequence, the AP frequency must be nonzero. We showed that GABA contributes to an increase in the leak conductance, which in turn flattens out the kink in the steady-state *I-V*-curve, and switches the excitability from class-I to II. Thus, heuristically, the critical leak conductance *ɡ*_l,0_ for which the kink disappears is

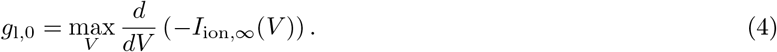

This condition, while sufficient for class-II excitability, is not necessary, however. At low firing rates, the positive feedback from a minor inflection in the *I*_∞_ (*V*) curve can be too weak to generate slow AP frequencies.

Nonetheless, our bifurcation analysis shows that (4) is one of the conditions for the system to be at the BTC point (Supplementary Text, Theorem 1), which localizes the switch in neuronal excitability to a single point in parameter space. Away from that BTC-point, the dynamical transitions in neuronal resonance, bistability, and the switch in excitability occur sequentially at three different points in the (*I*_e_,*ɡ*_l_)-plane. Depending on the precise neuron model, these transition points can be quite separated (Fig. 7, Supplementary Fig. S7i,iv).

### Universality and organization of transitions in AP dynamics across experiments and theory

As the BTC unfolding generically captures the modulation of AP dynamics, it provides an integrated explanation for a large number of related AP transition phenomena observed experimentally. Changing the neuron’s intrinsic conductance increases the non-zero onset AP frequency of fast spiking inter-neurons in rat somatosensory cortex (Tateno et al., 2004), and while pyramidal neurons are class-I excitable *in vitro*, they are class-II *in vivo* (Prescott et al., 2008b). Manipulating potassium conductances through cholinergic modulation of M-currents (Stiefel et al., 2008), changing the potassium channel density (Golomb et al., 1997; Zeberg et al., 2015) in pyramidal cells, or applying 4-AP to mesencephalic V neurons (Liu et al., 2008; Yang et al., 2009) are known to alter the excitability class of neurons. During neuronal development, conductance changes adjust AP latency and precision (Franzen et al., 2015).

Moreover, reexamining recordings of neurons in rat somatosensory cortex (Fig. 6A in Tateno et al. (2004)), of pyramidal and inter-neurons in prefrontal cortex (Fig. 1A in Fellous et al. (2001)) as well as stellate cells of the medial entorhinal cortex (Fig. 4-6 in Alonso & Klink (1993)) reveals signatures of conductance-induced stuttering or bistability, which were not explicitly the target of investigation in those studies. Additionally, resonances near threshold caused by peri-threshold or AP generating currents have been observed experimentally (Alonso & Llinas, 1989; Alonso & Klink, 1993; Gutfreund et al., 1995; Hu et al., 2002; Erchova et al., 2004). Increasing the outward potassium conductance in phasic vestibular neurons (Beraneck et al., 2007) or the background conductance in hippocampal neurons (Prescott et al., 2008b) are known to increase the resonance frequency. All of these experimental observations are consistent with the transition structure of the BTC-bifurcation point identified here.

Transitions in excitability and resonance have been studied intensively from a theoretical perspective, as the excitability class of neurons strongly influences the neuron’s selectivity to stimulus features (Rinzel & Ermentrout, 1989; Izhikevich, 2010; Robinson & Harsch, 2002; St-Hilaire & Longtin, 2004), the precision of spike timing (Gutkin & Ermentrout, 1998; Gutkin et al., 2005; Tateno & Robinson, 2006; Broiche et al., 2012), correlational coding (Hong et al., 2012), information processing and storage (Schleimer & Stemmler, 2009; Lengyel et al., 2005), as well as the collective dynamics on the network level (Hansel et al., 1995; Ermentrout, 1996; Ratté et al., 2013). The ordinary Bogdanov-Takens bifurcation has been characterized as an organizing center for transitions to resonance (Izhikevich, 2010; Ermentrout & Terman, 2010) and saddle-node-loop points to induce bistability (Izhikevich, 2010; Ermentrout & Terman, 2010). Potassium channel density can induce changes in excitability class as observed in simulations of fly neurons (Berger & Crook, 2015) and cortical neurons (Golomb et al., 1997; Arhem et al., 2006; Arhem & Blomberg, 2007; Zeberg et al., 2015). The BTC-structure captures these individual transitions in a single unifying framework.

In more complex neuron models, additional degenerate bifurcation points co-exist with the BTC point. For example, in the Connor-Stevens model neuron (Connor & Stevens, 1971; Connor et al., 1977), a nearby swallow-tail bifurcation induces two extra cusp points in the saddle-node bifurcation lines (not shown). Mixed-mode oscillations and transitions to chaos can also result in restricted regions around the double limit cycle bifurcation. Consistent with singular bifurcation theory (De Maesschalck & Wechselberger, 2015), the BTC structure, however, dominates the overall topology.

The theory developed here applies to non-bursting neurons. Computational models predict more complex transitions in the presence of three or more time scales (Drion et al., 2015) used to describe additional firing rate adaptation or bursting behavior. While some CA3 pyramidal cells fire APSs in bursts (Wong & Prince, 1981), and Hemond et al. (2008) classify one in five pyramidal cells as a burster, we did not encounter bursting cells in this study. Nevertheless, the theory may be extended to bursting neurons by mapping the slow dynamics onto trajectories within the BTC-structure that describes the faster APs (Osinga et al., 2012). Our approach to study AP modulation could be even further extended to higher degenerate bifurcation structures (Khibnik et al., 1998; Krauskopf & Osinga, 2016), which could provide global insight into the modulation of bursting dynamics (Franci et al., 2014).

### Control of neuronal excitability and network dynamics

How excitability and resonance at the single-cell level affect the population behavior of interconnected neurons has been extensively studied (e.g. Ermentrout (1996); Brown et al. (2004); Izhikevich (2010); Ermentrout & Terman (2010); Ratté et al. (2013)). Synchronization and information transmission between neurons depends on the functional form of the neuron’s phase response, which measure the shifts in the timing of A Ps due to oscillatory or pulsed stimuli (Ermentrout & Kopell, 1991; Ermentrout, 1996; Brown et al., 2004; Salinas & Sejnowski, 2001; Schleimer & Stemmler, 2009). Slow changes to a neuron’s intrinsic properties, for instance through spike frequency adaptation currents, will change the neurons’ phase response curves and hence network synchronization (Ermentrout et al., 2001). As a proof of principle, we used computational models of networks to show that also inhibitory pre-synptic neurons can provide sufficient tonic GABAergic drive to synchronize dynamically selected sub-sets of principal neurons. Comparatively little attention has been given to the role of dynamic bistability in controlling the number of synchronized neurons, even though the range of currents for which bistability exists can be large (Drion et al., 2015). These effects have potential roles in neuronal coding and information transmission, such as binding by synchrony (von der Malsburg, 1991; Singer & Gray, 1995) or communication through coherence (Fries, 2005) and can facilitate top-down processing across network layers (Engel et al., 2001; Lengyel et al., 2005), underscoring the importance of conductance and excitability for computation in the brain.

## Materials and Methods

### Slice preparation

Brain slices were prepared from Mongolian gerbils (Meriones uniguiculatus) of postnatal day (P) 10 to 18, as described in Ammer et al. (2012). In brief, animals were anesthetized, sacrificed, and then brains were removed in cold dissection solution containing (in mM) 50 sucrose, 25NaCl, 25NaHCO_3_, 2.5KCl, 1.25NaH_2_PO_4_, 3MgCl_2_, 0.1 CaCl_2_, 25 glucose, 0.4 ascorbic acid, 3 myo-inositol and 2Na-pyruvate (pH 7.4 when bubbled with 95% O_2_ and 5% CO_2_). Subsequently, 200 *μ*m thick transverse slices containing the DNLL (P10-11) or 300 *μ*m thick horizontal slices containing the hippocampus (P16-18) were taken with a VT1200S vibratome (Leica). Slices were incubated in extracellular recording solution (same as dissection solution but with 125 mM NaCl, no sucrose, 2mM CaCl_2_ and 1mM MgCl_2_) at 36°C for 45 minutes, bubbled with 5% CO_2_ and 95% O_2_.

### Electrophysiology

In the recording chamber, slices were visualized and imaged with a TILL Photonics system attached to a BX50WI (Olympus) microscope equipped with gradient contrast illumination (Luigs and Neumann). All recordings were carried out at near physiological temperature (34 - 36°C) in current-clamp mode using an EPC10/2 amplifier (HEKA Elektronik). Data were acquired at 50kHz and filtered at 3kHz. The bridge balance was set to 100% after estimation of the access resistance and was monitored repeatedly during recordings. The internal recording solution consisted of (in mM): 145K-gluconate, 5KCl, 15HEPES, 2 Mg-ATP, 2 K-ATP, 0.3 Na_2_-GTP, 7.5 Na_2_- Phospocreatine, 5K-EGTA (pH 7.2). 100 *μ*M Alexa 488 or 568 were added to the internal solution to control for cell type and location. To apply an artificial extrinsic leak conductance during recordings, an analogue conductance amplifier (SM-1, Cambridge Conductance) applied a constant conductance with a reversal potential equal to the neuron’s resting potential. GABA was puffed at an initial concentration of 500 *μ*M with continuous low pressure controlled by a picospritzer (Picospritzer III, Science Products). Glycinergic and glutamatergic synaptic inputs were blocked with 0.5 *μ*M Strychnine, 20 *μ*M DNQX, 10 *μ*M R-CPP, and GABAergic inputs were blocked with 10 *μ*M SR95531 in all experiments except when GABA was used as an agonist.

### Data Analysis

The intrinsic leak conductance was estimated from the voltage responses V (t) to small negative step currents of amplitude *δI*_e_ at *t* = 0 of 0.5 s duration introduced into the neuron. The average of 50 such traces was fitted by the voltage curve *V* (*t*) = *V*_min_ + *V*_0_ exp(- *t*/τ_L_). The leak was estimated as *ɡ*_L_ = *δI*_e_/(*V*_min_ − *V*_0_), and the capacitance as *c_m_* = τ_L_*ɡ*_L_. To validate the dynamic-clamp method, we determined the relationship between the imposed external leak *ɡ*_L,e_and the measured leak *ɡ*_L_. This relationship was linear with a slope very close to 1. The capacitance remained constant for different externally applied leak conductances (cf. Supplementary Fig. S2A,B). 1 s long depolarizing step currents were used to determine the *f-I*-curve and the area of bistability. The AP current threshold was estimated by a semi-automated search using 0.5 s long step currents. Voltage deflections crossing -20mV were classified as APs (Todd & Andrews, 1999). For the firing to be classified as periodic, APs had to occur throughout the 1s long stimulation at intervals with a coefficient of variation (CV) less than 50%. Spike frequency f was determined by the average of the inverse inter-spike intervals. If there were fewer than four APs, the inverse latency to the first AP was included in the average. A transition between bistable and periodic APs was deemed to occur when the CV of the inter-spike intervals changed by more than a factor of 1.5 between successive step currents. Clear alternations between periodic and non-periodic APs were seen in the voltage traces in the regimes of bistability. The spike onset frequency *f*_0_ was determined by fitting the *f-I*-curve to *f* (*I*_e_) = Θ (*I*_e_ − *I*_0_) [*a* (*I*_e_ − *I*_0_)^b^ + *f*_0_] with fit parameters *f*_0_ ≥ 0, *I*_0_, *a* ≥ 0, *b* and Θ a step-like threshold. Standard errors for the parameter estimates were used. The principal Floquet multipliers λ were estimated by fitting an exponential decay towards the steady state AP amplitude. Resonance frequencies were estimated by inducing a ZAP stimulus *I*_ZAP_ of the form

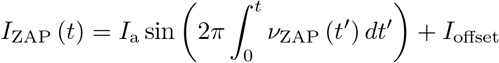

where the time dependent instantaneous frequency *ν*_ZAP_ (*t*) was ramped linearly from 0 to 25 Hz in 30 s and, for controls, from 25 Hz down to 0 Hz in 30 s. The ZAP current amplitude *I_a_* was calibrated before each sweep to yield a voltage deflection of ±5mV in the low frequency limit. The constant offset current *I*_offset_ was adjusted to make the cell be either just below or above spike threshold. The frequency-resolved impedance *Z* (*ν*) was estimated from the discrete Fourier transforms of *δI*(*t*) and *δV* (t), with 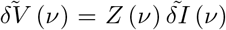. To estimate the resonance frequency, we fitted |*Z* (*ν*)|^2^ to the impedance of an RCL circuit of the form |*Z*_RCL_ (*ν*)|^2^ = (*a* + *bν*^2^) / (*ν*^4^ + *cν*^2^ + *d*), with fit parameters (*a,b,c, d*). Frequency components of 0.5Hz and less were dropped to exclude effects of slow drifts (Erchova et al., 2004). The resonance frequency *ν*_R_ was then determined by *ν*_R_ = argmax_*ν*≥0_ |*Z*_RCL_ (*ν*)|.

## Modeling

Conductance based models were constructed and simulated using a custom dynamical systems package we developed for Wolfram’s Mathematica 11.

**CA3 Pyramidal Neuron Models** A single compartment conductance-based neuron model based on a simplified regular firing CA3 neuron model by Migliore et al. (2010) was fit to the experimentally observed spike responses, AP shapes and phase-plane dynamics manually. The voltage *V* of the model evolved in time *t* according

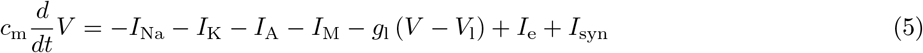

with specific capacitance 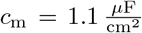, leak conductance 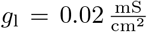 and reversal potential *V*_l_ = -60 mV, *I*_e_ an externally applied and *I*_syn_ the synaptic trans-membrane current. A minimal model that fits the data includes four active currents: A fast sodium *I*_Na_ = *ɡ*_Na_*m*^3^*h* (*V* − *V*_Na_), a potassium delayed rectifier *I*_K_ = *ɡ*_K_*n*^4^ (*V* − *V*_K_), a transient A-type *I*_A_ = *ɡ*_A_*n*_A_*l*_A_ (*V* -*V*_K_), and an M-type *I*_M_ = *ɡ*_M_*m*_M_ (*V* − *V*_K_) adaptation current with maximal specific conductances 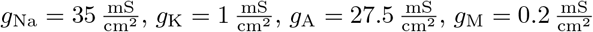 and reversal potentials *V*_Na_ = 50 mV, *V*_K_ = − 90mV. The activation and inactivation variables *a* ∊ {*m,h,n,n*_A_,*l*_A_,*m*_M_} evolve as in Migliore et al. (2010) but were shifted in voltage to match the data. Their dynamics are given by

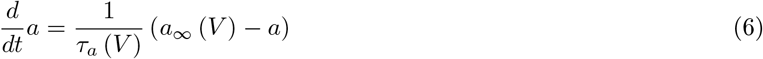

with

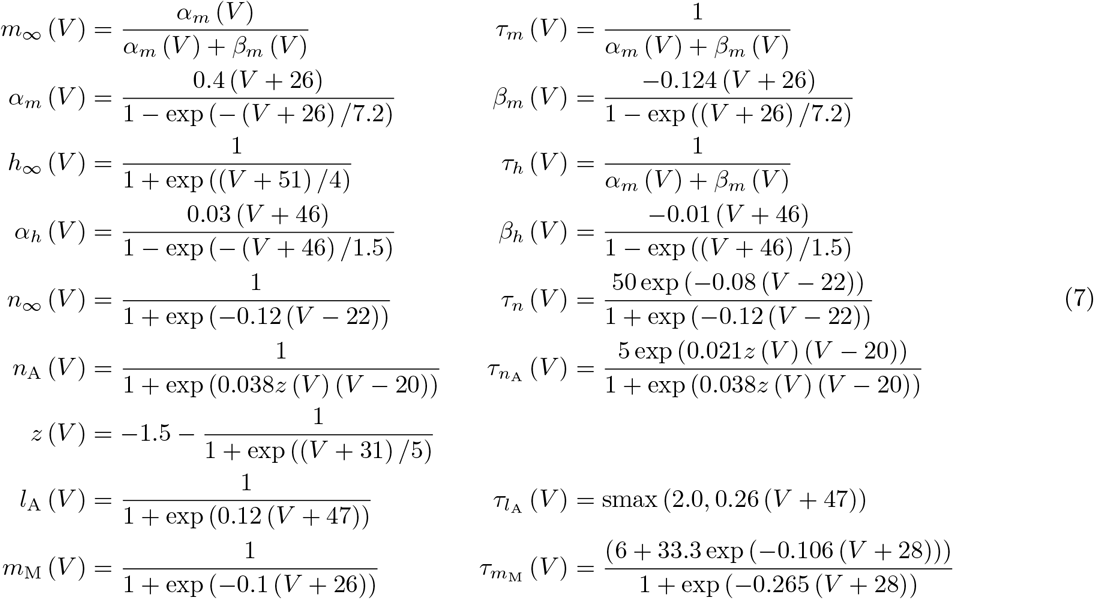

where for 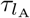 we used a soft-max function smax (*x, y*) = log (exp (*kx*) + exp (*ky*))/*k* with *k* = 40 to ensure continuity of the derivatives and enable numerical continuation. In Fig. 3 we used a surface area of *a* = 0.0001 cm^2^ to match the experimentally observed absolute conductances.

**Other Neuron Models** To show the generality of our theory we also analyzed the Wang-Buzsaki inter-neuron model (Wang & Buzsáki, 1996), the generic Morris-Lecar neuron model (Morris & Lecar, 1981) and the fast spiking neuron by Erisir et al. (1999), the Rinzel model (Rinzel, 1985), Rose-Hindmarsh model (Rose & Hindmarsh, 1989), Connor-Stevens model (Connor & Stevens, 1971), Golomb model (Golomb et al., 2007) and reduced pyramidal neuron models (Prescott et al., 2008b; Traub & Miles, 1991; Ermentrout & Kopell, 1998) using the parameter values as in the references. In the Wang-Buzsaki model the capacitance was set to 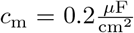 in Fig. 7D,E and to 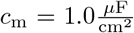 otherwise. For convenience, the full equations of all models used are listed in the Supplementary Methods section.

## Bifurcation Analysis

Bifurcation diagrams and Floquet multipliers (Cesari, 1971) were numerically computed using the continuation software AUTO (Doedel et al., 2007) interfaced with our dynamical systems package that automatically generates c++ code used in AUTO from the model definitions.

## Resonance Detection

Neuronal resonance frequencies were determined from the linear response to sinusoidal input stimuli around a steady state *x*_0_ = (*V*_0_, *a*_2,0_,…, *a*_*N*,0_)^T^ using equation (8). The impedance is

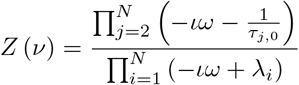

where 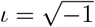 and λ_*i*_ are the eigenvalues of the Jacobian *Df* (*x*_0_) (see Supplementary Methods). The transition to resonance was detected by numerical continuation of the condition 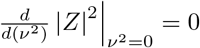 using AUTO. Saddle-point to focus transitions were detected by continuation of the zeros of the discriminant for the characteristic polynomial det (*Df* (*x*_0_) − λ). Resonance frequencies *ν*_R_ where determined by *ν*_r_ = argmax_*ν*≥0_ |*Z* (*ν*)|.

### Mathematical Analysis: Organization of Neuronal Excitability and Resonance Transitions

The impact of leak currents on neuronal excitability was mathematically studied using conductance-based neuron models of the form

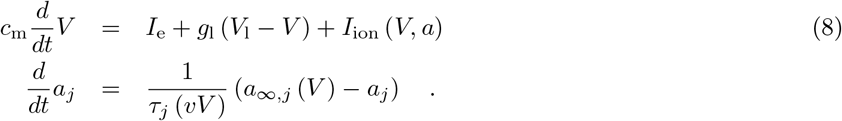

The active ion currents are assumed to have the general from

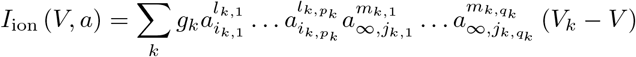

with maximal conductances *ɡ*_k_, reversal potentials *V*_k_, time constants τ_*j*_, and allowing for an arbitrary number of activation and inactivation variables *a* = (*a*_1_, *a*_2_,…, *a_N_*) and possible substitutions with steady state activations *a_∞,j_*(*V*) each with their own real non-negative exponents *l_k,p_*,*m_k,q_* ≥ 0. For simplicity of the derivations we use a non-standard sign convention for *I*_ion_ (*V*, *a*) treating inward, depolarizing currents as positive.

By using a combination of a center-manifold and normal form reduction (Kuznetsov, 2005) together with multiple bifurcation theory (Guckenheimer, 1986) we prove the following results:

#### Theorem

Every neuron with class-I excitability of the form (8) has a Bogdanov-Takens-Cusp point (Dumortier et al., 1991) in the parameter space of input, leak conductance, and capacitance (*I*_e_,*ɡ*_l_,*c*_m_).

To proof this we make use of three observations: first, the bifurcation parameters *I*_e_, *ɡ*_l_, and *c_m_* are general parameters occurring in all conductance-based neuron models and only appear as coefficients of constant or linear terms in *V* in the dynamical equations. Second, the dynamics of the gating variables are coupled solely through the membrane potential *V*. holds. Third, a positive feed back on arbitrarily slow times scales generating class-I excitability in (??) is possible if there is some voltage *V*^+^ such that the *f-I*-curve (3) has positive slope, i.e. 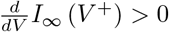.

By using these properties we can adjust the three parameters (*I*_e_,*ɡ*_l_,*c*_m_) and the state vector (*V, a*) such that the system is at a co-dimension three BTC point (Dumortier et al., 1991). The proof first expresses all parameters and state variables as functions of V alone to satisfy *N* + 2 out of the *N* + 3 constraints for a codimension-3 BTC bifurcation (Kuznetsov, 2005). The last constraint reduces to an equation for *V* that be can be shown to have a solution. A detailed mathematical proof is given in the Supplementary Text.

#### Proposition.

If

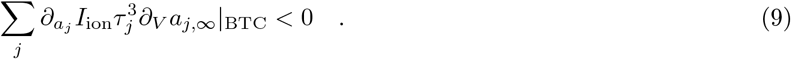

the BTC point is of focus or elliptic type.

The proof follows by showing that the condition (9) entails the conditions for a BTC point to be of focus and elliptic type (Dumortier et al., 1991) (see Supplementary Text for details).

In appropriate state coordinates *u*_1_,*u*_2_ the unfolding of the BTC point takes the form

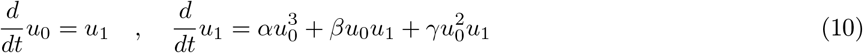

that represents the simplest neuron model that in dependence of parameter *α, β* and *γ* captures all the dynamical regimes observed in the neuronal excitability transitions. For a neuron model satisfying the conditions of the above theorem a parameter dependent coordinate transformation exists (Dumortier et al., 1991) that up to higher order terms transforms it into the equations (10) near the BTC point (see Supplementary Text for details).

### Network and Resonance Simulations

Networks of *N* generic conductance based class-I model neurons (Wang & Buzsáki, 1996) of the form (8) using original parameter values and with 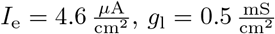 and *V*_l_ = − 60 mV (cf. Fig. 6) were coupled all-to-all via excitatory synapses. A synapse from pre-synaptic neuron *j* added a current

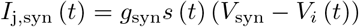

to post-synaptic neuron *i* with membrane potential *V_i_*. For weak (strong) excitatory synapses we used *ɡ*_l_ = 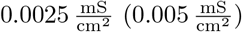, *V*_syn_ = 0mV. The activation *s* (*t*) evolved as

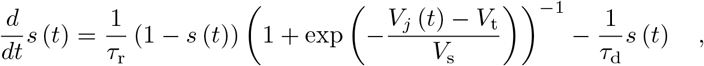

with τ_r_ = 0 ms, τ_d_ = 1.5 ms, *V*_t_ = − 20 mV, *V*_s_ = 1mV. Activation of inhibitory neurons was simulated by Poisson spike trains activating shunting synapses with τ_d_ = 5 ms and *V*_syn_ = − 51.4 mV, resulting in a weakly fluctuating shunting conductance around 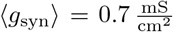, sufficient to shift neurons to class-II excitability. Synchrony was measured by the vector strength 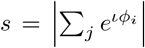 with *ϕ*_*i*_ the instantaneous phase of neuron *i* estimated by linear interpolation from 0 to 2π between spikes. Spikes were determined by positive voltage crossings at *V*_t_ = − 20 mV.

## Author Contributions

C.K. and M.S. designed research; C.K. derived the theoretical results and carried out numerical analysis with input from M.S.; all authors designed experiments; J.A. performed experiments; C.K. analyzed the data; C.K. and M.S. wrote the manuscript with input from J.A. and A.H.

## Acknowledgements

We thank J. Hudspeth, C. Bargmann, E. Siggia, M. Gutnick, L. Paninski, T. Geisel, and A. Neef for helpful discussions. This work has been supported by the German Federal Ministry for Education and Research through the Bernstein Center for Computational Neuroscience Munich (Grant 01GQ0440), and fellowships from the Rockefeller University and from the Kavli Foundation to CK.

